# A Complex Interplay of Anionic Phospholipid Binding Regulates 3’-Phosphoinositide-Dependent-Kinase-1 Homodimer Activation

**DOI:** 10.1101/566430

**Authors:** Gloria de las Heras-Martínez, Véronique Calleja, Remy Bailly, Jean Dessolin, Banafshé Larijani, Jose Requejo-Isidro

**Affiliations:** Instituto Biofisika (CSIC, UPV/EHU), 48490 Leioa, Spain; Protein Phosphorylation Laboratory, The Francis Crick Institute, 1 Midland Road, NW1 1AT, London, UK; Institute of Chemistry & Biology of Membranes & Nanoobjects (UMR 5248 CBMN) CNRS – Université de Bordeaux - Bordeaux INP All. Geoffroy Saint-Hilaire, 33600 Pessac, France; Cell Biophysics Laboratory, Ikerbasque Basque Foundation for Science, Instituto Biofisika (CSIC, UPV/EHU) & Research Centre for Experimental Marine Biology and Biotechnology (PiE), University of the Basque Country (UPV/EHU), Leioa 48940, Spain; Centro Nacional de Biotecnología (CSIC). Darwin, 3, E28049 Madrid, Spain

## Abstract

3’-Phosphoinositide-dependent-Kinase-1 is a master regulator whereby its PI3-kinase-dependent dysregulation in human pathologies is well documented. Understanding the direct role for PtdIns(3,4,5)P_3_ and other anionic phospholipids in the regulation of PDK1 conformational dynamics and its downstream activation remains incomplete.

Using advanced quantitative-time-resolved imaging, FCS and molecular modelling, we show an interplay of antagonistic binding effects of PtdIns(3,4,5)P_3_ and other anionic phospholipids, regulating activated PDK1 homodimers. We demonstrate that phosphatidylserine maintains PDK1 in an inactive conformation. The dysregulation of the PI3K pathway affects the spatio-temporal and conformational dynamics of PDK1 and the activation of its downstream substrates.

We establish an anionic-phospholipid-dependent model for PDK1 regulation, depicting the conformational dynamics of multiple homodimer states. The dysregulation of the PI3K pathway perturbs equilibrium between the PDK1 homodimer conformations. Our findings indicate that the alteration of specific basic residues of PDK1-PH domain leads to its constitutive activation, a potential significance in different types of carcinomas.

## Introduction

PDK1 propagates extracellular signals downstream of Phosphoinositide 3-kinase (PI3K) through the phosphorylation of a plethora of AGC kinases involved in growth, survival and cell motility [1, 2]. PDK1 has many kinase-dependent and independent functions, and fulfils the criteria of being a master regulator of proliferative signalling. The physiological evidence is clear as alteration in PDK1 expression or activity have been found many types of cancer [3]. PDK1 can associate with the plasma membrane (PM) by interaction with phosphatidylinositol (3,4,5) trisphosphate (PtdIns(3,4,5)P_3_), phosphatidylinositol (4,5) bisphosphate (PtdIns(4,5)P_2_) and phosphatidylserine (PtdSer) via its pleckstrin homology (PH) domain [4-6], and through other adaptor proteins such as Grb14 [7], Grp78 [8] and Freud-1/Aki1 [9]. Despite being mainly localised in the cytoplasm and at the PM, pools of PDK1 have been detected in the nucleoplasm where many of its substrates also reside [10-13]. The high-affinity specific interaction of PDK1 with PtdIns(3,4,5)P_3_ through its K465 residue on its PH domain has been well established [5, 14, 15]. However, reports on whether PtdIns(3,4,5)P_3_ is solely responsible for PDK1 recruitment to the PM are contradictory [4, 6, 16], suggesting a complex interplay between PDK1 binding to phosphoinositides and other anionic phospholipids. PtdSer is the predominant anionic species and at higher concentrations than the phosphoinositides at the inner leaflet of the plasma membrane [17]. PtdSer has been reported to be involved in the membrane localisation of PDK1 through its basic residues R466 and K467 [6] but the role of this interaction has not yet been fully elucidated. It has been shown that PDK1 is constitutively active since it is phosphorylated on its activation loop at S241 and is active *in vitro*, [18]. Under these circumstances when PDK1 localised to the plasma membrane it phosphorylated PKB/Akt. However, it was also shown *in situ* that PDK1 itself could be regulated upon stimulation [19, 20]. More recently, PtdIns(3,4,5)P_3_ binding to PDK1 in live cells was shown to elicit the formation of PDK1 homodimers [13, 19] and to trigger the autophosphorylation of the PDK1 PH domain residue 513. This, was suggested to be due to the disruption of an autoinhibitory PDK1 homodimer conformer [13, 21, 22]. This mechanism still remains to be defined. Altogether, these findings pointed towards an elaborate fine-tuning of PDK1 conformational dynamics upon stimulation in cells, that involve, the interaction with negatively charged lipids such as PtdIns(3,4,5)P_3_, PtdIns(4,5)P_2_ and PtdSer, autophosphorylation of T513 and homodimerisation. However, our understanding of the interplay between these events and the associated regulation that enables the phosphorylation of PDK1 substrates remains largely unknown. Addressing these issues has been hampered by the lack of suitable *in situ* quantitative methods for interrogating the spatio-temporal sequence of protein-lipid interactions. We have recently reported the development of a precise and well-resolved quantitative imaging method based on fluorescence lifetime imaging (FLIM) that overcomes these technical obstacles [23]. Here, we were able to monitor *in situ* the anionic phospholipid-mediated regulation of PDK1 homodimerisation and localisation leading to PDK1 activation and how this was linked to T513 autophosphorylation in cells with a normal or a dysregulated PI3K pathway. Using this methodology, we unravelled three major mechanisms for PDK1 regulation. Firstly, that PDK1 homodimerises in an activated conformation phosphorylated on T513 and capable of activating downstream substrates like PKB/Akt and SGK1. Secondly, that this homodimer formation was triggered by two opposite mechanisms: (i) The binding to PtdIns(3,4,5)P_3_ upon growth factor stimulation (ii) the loss of plasma membrane binding to other anionic phospholipids. This suggested a competitive regulatory mechanism involving PtdIns(3,4,5)P_3_ and other anionic phospholipids having opposite effects on PDK1 activation. Thirdly, we demonstrated that PDK1 activation through recruitment to the PM was not only dependent on its PH domain but its kinase domain played a prominent role in this process. The role of the kinase domain interaction was of particular importance in cells with a dysregulated PI3K pathway and an aberrant regulation of the PtdIns(3,4,5)P_3_ levels, leading to an abnormal PDK1 activation.

## Results

### Characterisation of anionic lipids binding to PDK1 PH domain *in vitro*

The nature of PDK1 interactions and regulation by anionic phospholipids at the plasma membrane, remains a subject of discussion. To understand the mechanism of interplay of the anionic phospholipids associations with PDK1, we characterised their *in vitro* interactions using fluorescently tagged PH domains of PDK1 (PH^PDK1^). PH^PDK1^ wild type and mutants were tested using a protein-lipid overlay assay, and their diffusional behaviour was measured by single-focus scanning Fluorescence Correlation Spectroscopy (FCS) [24] (Fig. 1). The positively charged amino acid site 465, previously associated with PtdIns(3,4,5)P_3_, and 466/467 associated with PtdIns(4,5)P_2_ and PtdSer binding [5, 6], were mutated to the neutral residues alanine (K465A and R466A/K467A) (Fig. S1A). We first determined PH^PDK1^ affinity for PtdIns(3,4,5)P_3_, PtdIns(4,5)P_2_ and PtdSer using a protein-lipid overlay assay. We confirmed that wild type-PH^PDK1^ bound PtdIns(3,4,5)P_3_ with a high affinity and PtdIns(4,5)P_2_ to a lesser extent (Fig. 1A). In line with previous reports detectable interactions with PtdIns(3,4,5)P_3_ or PtdIns(4,5)P_2_ were not observed with the K465A-PH^PDK1^ mutant even during extended incubations [5]. The R466A/K467A-PH^PDK1^ mutant however selectively lost its binding to PtdIns(4,5)P_2_ but not to PtdIns(3,4,5)P_3_.

**Fig. 1.**
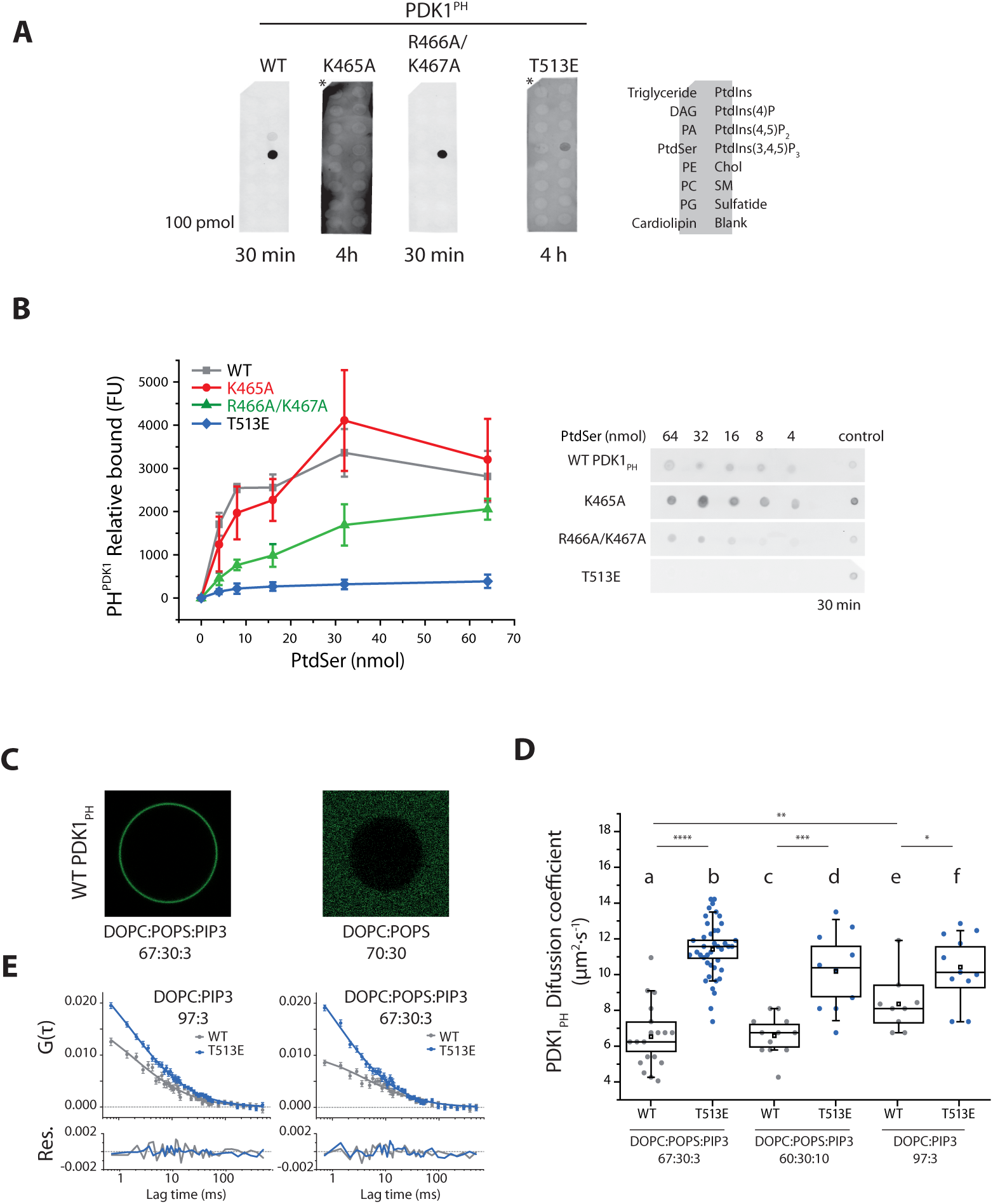
Characterisation of anionic lipids binding to PDK1 PH domain *in vitro*. **(A)**. Lipid protein-overlay assays showing the binding of WT-PH^PDK1^ and mutants to a selection of lipids spotted on nitrocellulose. **(B)** PH^PDK1^ wild type and mutants’ affinity for PtdSer. The relative binding of WT-PH^PDK1^ and mutants is represented on the graph. **(C)** Images of WT-PH^PDK1^ interaction to GUVs of different compositions. **(D)** WT-PH^PDK1^ and mutant T513E-PH^PDK1^ diffusion on GUVs of varied composition. T513E-PH^PDK1^ diffusion is at the detection limit of our system, indicating non-specific binding. **(E)** Representative FCS correlation curves, each corresponds to one GUV. N>8. Box: 2xSEM (95.4% confidence); Whiskers: 80% population. Mann-Whitney test *p<0.05.

Since the residues R466/K467 were previously proposed to be necessary for PDK1 interaction with PtdSer [6] we investigated the binding properties of the different mutants with this phospholipid. PtdSer binding to PH^PDK1^ was only detected after a 40-fold increase of the amount of lipid (from 100 pmol in Fig. 1A to 4 nmol in Fig. 1B), demonstrating that the *in vitro* PH^PDK1^ affinity for PtdSer was significantly lower than PtdIns(3,4,5)P_3_. The mutation K465A did not affect the binding to PtdSer, while the mutation R466A/K467A showed a four-fold reduction in affinity. These data suggested that despite the close proximity of these three residues (K465, R466A and K467A) on the PDK1 PH domain, their different binding properties enabled to discriminate the effect of PtdIns(3,4,5)P_3_ from the other anionic phospholipids.

We next determined whether post-translational modifications of the PH domain could also regulate PDK1 binding to the plasma membrane by affecting its association with anionic phospholipids. We and others had previously shown that PDK1 activation was dependent on the autophosphorylation of the PH domain residue the T513 upon PM translocation [13, 21, 25]. Therefore, we examined whether T513 phosphorylation affected the binding of PH^PDK1^ to anionic phospholipids. To this aim we prepared a PH^PDK1^ mutant that mimicked PH^PDK1^ phosphorylation on T513 (T513E-PH^PDK1^) [25]. Fig. 1A shows that even though T513E-PH^PDK1^ retained some ability to bind PtdIns(3,4,5)P_3_ it required an extended incubation time to be detected, reflecting a lower affinity for this lipid compared to the wild type species. In addition, T513E-PH^PDK1^ did not bind PtdSer even at the highest concentrations (Fig. 1B). These experiments suggested that the autophosphorylation of the PDK1 PH domain residue 513 upon stimulation could trigger a loss of affinity of the activated PDK1 for the PM.

To further understand the nature of the PDK1 – anionic lipid interaction using scanning FCS, we studied PH^PDK1^ diffusion on giant unilamellar vesicles (GUV) composed of DOPC, PtdIns(3,4,5)P_3_ and PtdSer mixed at varying proportions (Fig. S2). Diffusion of a protein on a lipid bilayer depends largely on the lipid-protein electrostatic interactions. A weak interaction of a protein will result on the protein “sliding on” the vesicle surface faster than the lipids in the mixture, its diffusion subjected only to the electrostatic attraction between the positive surface of the protein and the negatively charged surface of the vesicle. Modulation of the vesicle surface potential will, in turn, affect the diffusion coefficient of the protein. However, a protein tightly bound to a single lipid through a stereospecific interaction will diffuse at the same rate as the latter, irrespective of the surface potential of the vesicle [26].

Wild type PH^PDK1^ readily bound the outside layer of a GUV containing only 3% PtdIns(3,4,5)P_3_ (DOPC:DOPS:PIP3 (67:30:3)), but binding was not detected in DOPC:DOPS (70:30) GUVs when PtdIns(3,4,5)P_3_ was absent in the mixture (Fig. 1C). This behaviour reflected the results obtained from the protein-lipid overlay assay.

The diffusion coefficient of WT-PH^PDK1^ did not change in DOPC:DOPS containing GUVs when the concentration of the polyvalent, highly negative, PtdIns(3,4,5)P_3_ was changed from 3% to 10% (Fig 1D a, c). This confirmed that PH^PDK1^ bound PtdIns(3,4,5)P_3_ in a lipid-specific manner, rather than by a non-specific, charge-driven, interaction. Given the weak binding of PH^PDK1^ to PtdSer compared to PtdIns(3,4,5)P_3_ (Fig. 1B), and that binding of PH^PDK1^ to DOPC:POPS (70:30) GUVs was not observed, we did not anticipate any effect of PtdSer in the diffusion of PH^PDK1^. We found however that PH^PDK1^ diffused faster on GUVs containing PtdIns(3,4,5)P_3_ but lacking PtdSer (DOPC:PIP3 97:3) (Fig. 1D a and e). This indicated that PtdSer was able to bind to PH^PDK1^ only in combination with PtdIns(3,4,5)P_3_-binding. This behaviour could be explained by the close proximity of PH^PDK1^ to PtdSer upon binding to PtdIns(3,4,5)P_3_ GUV. This dual binding maybe due to either the binding of the 2 anionic phospholipids to the same PH domain or to the formation of a dimeric conformation where each lipid would bind to one PH. FCS alone would not be able to distinguish between a monomeric PH conformer versus a dimeric PH conformer. The phosphomimetic T513E-PH^PDK1^ mutant diffused in a similar manner in GUVs with different lipid compositions and it had a faster diffusion than WT-PH^PDK1^ in DOPC:PIP3 GUVs (Fig. 1D b, d and f). Since the affinity of T513E-PH^PDK1^ for PtdIns(3,4,5)P_3_ was very low (and negligible for PtdSer), this behaviour indicated the predominance of non-specific, electrostatically driven attraction of PH^PDK1^ positive patches to the negatively charged vesicle surface.

### Opposing effects of anionic phospholipids regulate the formation of activated PDK1 homodimers

Having established the *in vitro* binding properties of the different PDK1 phospholipids mutants and determined the effect of PtdSer on the binding of PDK1 PH domain to PtdIns(3,4,5)P_3_, we examined the interplay of these lipids in the regulation of full-length PDK1 in cells. To this end, we analysed the activation potential of wild type PDK1 and relevant PDK1 mutants through the study of their localisation and conformational dynamics. We refer to an activated PDK1 conformer the molecular conformation that enables the phosphorylation of its downstream targets.

Previously, we observed PDK1 homodimerisation in response to growth factor stimulation [13]. The interacting population of PDK1 had however not been directly determined but retrospectively calculated. Thus, in this investigation we directly quantified the interacting population of full-length protein PDK1 homodimers *in situ* using quantitative FRET-FLIM (Supporting Methods and Fig. S3). We also utilised automated image segmentation methods on intensity and FRET-FLIM images. The segmentation determined the spatial localisation of PDK1 in the subcellular compartments as well as the localisation of PDK1 homodimers in response to growth factor stimulation. FRET detected by FLIM as the energy transfer from ectopically expressed eGFP-myc-PDK1 (donor) to HA-PDK1-mCherry (acceptor) (Fig. S1B). Here, based on lifetime-analysis we determined an effective fraction of donor undergoing FRET, 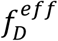, for every cell (Fig. S3A-B) and defined the “Dimerisation Efficiency” (*E_D_*) [23]. As opposed to the FRET efficiency, which was dependent on the relative abundance of interacting partners, *E*_D_ directly accounts for cell-to-cell variations in donor and acceptor concentration (Fig. S3C-E).

Initially, we analysed GFP-PDK1 and PDK1-mCherry recruitment profiles in NIH3T3 cells irrespective of their dimerisation status (Fig. 2A). To measure PDK1 recruitment levels at the plasma membrane we used an operator-free algorithm to segment the intensity images of the plasma membrane, cytoplasm and nucleus (Fig. S3F and S4). Intensity segmentation results indicated a predominantly cytoplasmic distribution of GFP-PDK1 and PDK1-mCherry in resting NIH3T3 cells (Fig. 2A). PDK1 translocation to the plasma membrane following an increase of PtdIns(3,4,5)P_3_ upon growth-factor stimulation, was prevented by the PtdIns(3,4,5)P_3_-binding site mutation K465A (Fig. 2A, b). In all cases, PDK1 recruitment to the NIH3T3 plasma membrane was compromised when PI3K was inhibited by LY294002 prior to stimulation. Nevertheless, despite the weak affinity of the isolated PH domain mutant T513E for PtdIns(3,4,5)P_3_ and other phospholipids *in vitro* (Fig. 1), full-length PDK1-T513E recruited to the PM upon PDGF stimulation. This result suggested a potential role for PDK1 kinase domain, either directly binding to the PM or by an allosteric effect on the PH domain affinity for the PM (Fig. 2, g).

**Fig.2.**
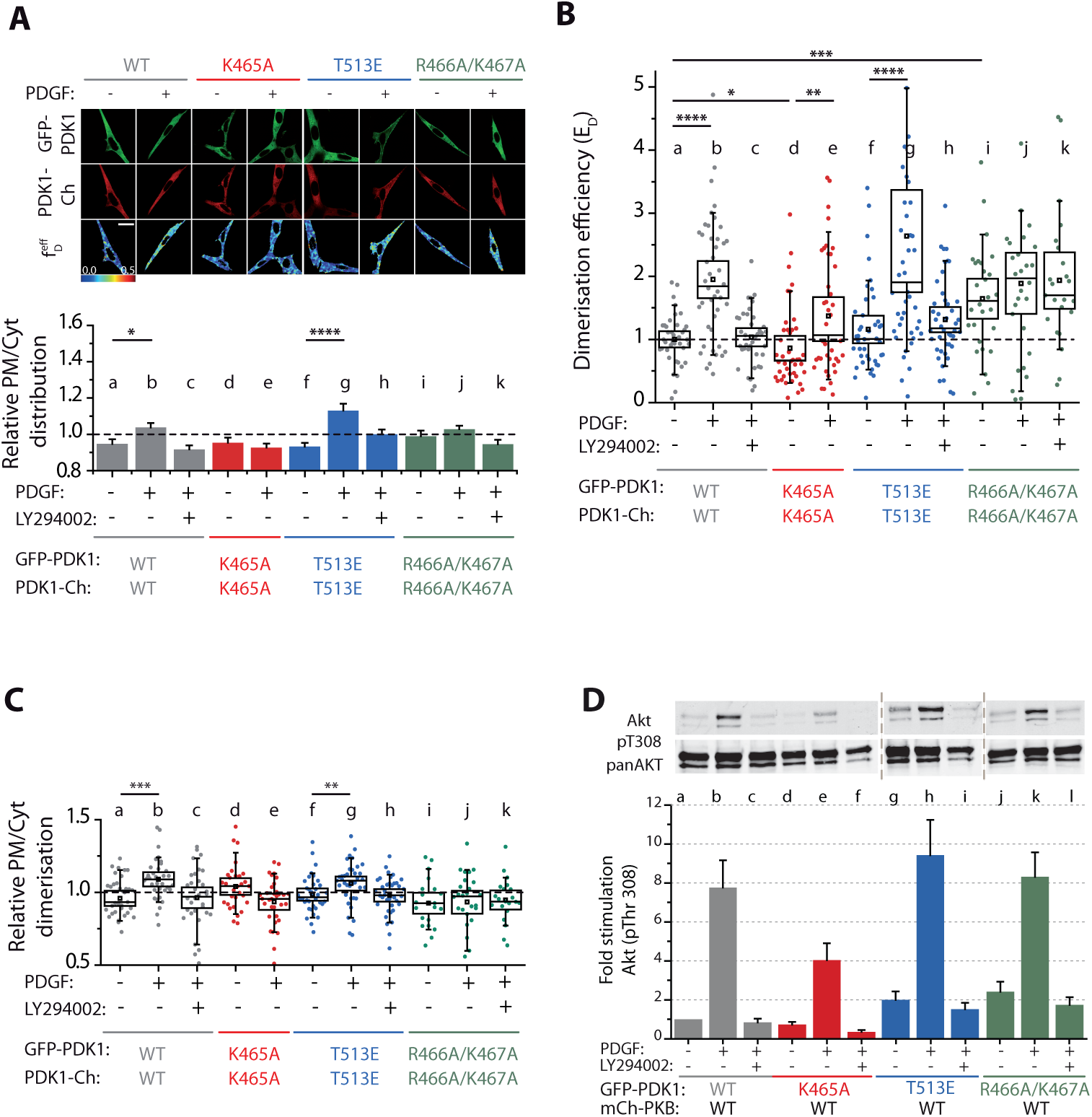
Opposing effects of anionic phospholipids regulate the formation of active PDK1 homodimers. All experiments were performed under resting conditions (-), stimulation of the PI3K pathway with PDGF (+) or treated with the PI3K inhibitor (LY294002) prior to stimulation. NIH3T3 cells were cotransfected with GFP-PDK1 and PDK1-mCherry **(A)** Representative intensity and FRET-FLIM *f*_*D*_ images of NIH3T3 cells cotransfected with GFP-PDK1 and PDK1-mCherry. The graph below represents the relative distribution of the fluorescent species at the PM and the cytoplasm of the cell segmented using an automated segmentation algorism. **(B)** Quantification of the homodimerisation efficiency of PDK1 wild type and mutants. *E* D calculation is based on FRET-FLIM measurements on a single-cell level, as illustrated in (A). **(C)** Localisation of the PDK1 homodimers in cell. The box and whiskers represent the relative amount of the dimeric species at the PM and in the cytoplasm. **(D)** Phosphorylation of Akt/PKB at T308 in NIH3T3 cells. Upper panel shows a representative western blot of phosphorylated Akt/PKB on T308 and of total Akt/PKB (pan AKT). The graph represents the ratio of T308 phosphorylation over total protein in each condition. The data were normalised to the condition basal wild type PDK1. Scale bar: 20 μm. (B) N>30. Error bars: SEM. (C) and (D) N>30. Box: 2xSEM (95.4% confidence); Whiskers: 80% population. Mann-Whitney test *p<0.05. Three independent experiments.

The mutation of the PDK1 PH domain residues R466A/K467A was previously described to impair the binding of PDK1 to PtdSer and the localisation at the PM of NIH3T3 cells [6]. Our *in vitro* experiments showed that in addition to PtdSer, the mutation also precluded the binding to PtdIns(4,5)P_2_ (Fig. 1 A). Nonetheless the overall localisation of full-length GFP-PDK1 and PDK1-mCherry in NIH3T3 cells was not affected by the R466A/K467A mutation.

Next, we monitored the homodimerisation of PDK1 by acquiring, *in situ*, full-length GFP-PDK1 and PDK1-mCherry wild type and mutants homodimerisation by FRET-FLIM. We observed that dimer formation and dimer localisations were not systematically following the localisation pattern determined for the total amount of protein, described above (observations for total protein, based on intensity measurements, did not take into account whether the protein was in a dimer form). The *E*_D_ values for WT PDK1 showed an increase of PDK1 homodimerisation in response to growth factor (PDGF) stimulation, while it remained at basal level when PI3K was inhibited by LY294002 prior to stimulation (Fig. 2B, a-c). Consistent with the latter, the loss of PtdIns(3,4,5)P_3_ binding (K465A) precluded the strong dimerisation of PDK1 upon PDGF stimulation. A small increase in *E*_*D*_ was still detected possibly due to unspecific electrostatic interactions (Fig. 2B, d-e).

Mimicking PDK1 phosphorylation with the PH domain mutation T513E had been shown to increase PDK1 activity [13, 25]. Additionally, residue 513 phosphorylation was known to relieve a PDK1 autoinhibitory conformation [21]. This led to the hypothesis that T513 phosphorylation induced a disruption of PDK1 autoinhibited homodimers and resulted in the formation of active PDK1 monomer [13, 21]. However, by determining the E_D_ we illustrate here that T513E mutation did not prevent PDK1 homodimerisation Figure 2B, and that the dimer regulation was similar to the WT-PDK1 (Fig. 2B f-h). This demonstrated that the activation of PDK1 by phosphorylation of 513 did not induce PDK1 monomers but resulted in an enhancement, of PDK1 homodimer population.

The loss of PtdSer and PtdIns(4,5)P_2_ binding through the R466A/K467A mutation did not prevent PDK1 homodimerisation. On the contrary, the population of R466A/K467A-PDK1 dimers was significantly higher than the WT-PDK1 in basal conditions and did not change upon either growth factor stimulation or PI3K inhibition (Fig. 2B, i-k). Since the constitutive formation of the R466A/K467A-PDK1 homodimers was independent of the modulation of PI3K, these data suggested that R466A/K467A-PDK1 homodimer formation, unlike WT-PDK1, occurred through a mechanism independent of its binding to PtdIns(3,4,5)P_3_ and other anionic phospholipids.

To understand the mechanisms of PDK1 homodimer formation, we specifically analysed their subcellular localisation. Using image segmentation of the FRET-FLIM images (Fig. 2C) differences were revealed between the localisation of total PDK1 (irrespective of its homodimerisation status, Fig. 2A) and the population of PDK1 homodimer species. Segmentation of FRET-FLIM images demonstrated that WT-PDK1 dimers were recruited to the plasma membrane in NIH3T3, upon the increase of PtdIns(3,4,5)P_3_ levels (Fig. 2C, a-c). The localisation of the PDK1 mutants that did not bind PtdIns(3,4,5)P_3_ via their PH domain (K465A), were still retained at the PM in basal conditions, indicating that the localisation of the PDK1 homodimers was due to PM interactions involving other factors than the binding of the PH domain to PtdIns(3,4,5)P_3_. The localisation of the T513E mutant followed the pattern of the wild type protein, suggesting that the mechanisms involved in the formation of the WT and T513E-PDK1 homodimers were similar. The R466A/K467A-PDK1 dimers did not localise to the plasma membrane even after growth factor stimulation but they predominantly localised in the cytoplasm. Therefore, the R466A/K467A-PDK1 mutant behaviour suggests a different mechanism for the formation of PDK1 homodimers.

Our results demonstrated that the increased formation of PDK1 homodimers was not only the result of growth factor stimulation, but could also occur in a constitutive manner due to mutation. We therefore determined the activation status of WT-PDK1 and its mutants. Our results also showed that the activating mutant T513E-PDK1 was also a homodimer, raising the possibility that the homodimer conformer of PDK1 would be activated and capable of phosphorylating downstream targets. Therefore, we monitored the effect of the mutants on the phosphorylation of Akt/PKB, a PDK1 substrate. Fig. 2 shows the direct activation of Akt/PKB by PDK1 reported by the phosphorylation of the T308 residue in Akt/PKB’s activation loop. Phosphorylation of Akt/PKB by WT-PDK1 increased upon PDGF stimulation (shown in grey). This increase correlated with the augmentation of WT-PDK1 homodimers and their localisation at the PM (Fig. 2B, b and 2C, b). PDK1 PtdIns(3,4,5)P_3_-binding defective mutant K465A, moderated T308 phosphorylation increase upon stimulation, but did not abolish it completely (shown in red). This also correlated with K465A-PDK1 basal localisation at the PM and residual formation of these mutant homodimers upon stimulation (Fig. 2C, d and 2B, e). Both the activating T513E-PDK1 and the PM localisation-defective R466A/K467A-PDK1 triggered an increase in T308 phosphorylation, upon stimulation, that was similar to the WT-PDK1 homodimers (in blue and green in Fig. 2D, respectively). The common aspect of the WT-PDK1, T513E and R466A/K467A was an elevated homodimer formation, despite their different localisation patterns (Fig. 2B). It is important to note that irrespective of PDK1 homodimerisation status, Akt/PKB requires prior binding to PtdIns(3,4,5)P_3_ before being phosphorylated by PDK1 at T308 [10]. Therefore, in resting conditions or after PI3K inhibition, Akt/PKB phosphorylation was reduced despite the increased levels of PDK1 homodimers (Fig. 2Dj, l and 2Bi, k). Both T513E and R466A/K467A mutations induced a slightly higher basal phosphorylation of Akt/PKB compared to WT. Although this basal increase in Akt/PKB phosphorylation was already described for the activating T513E mutant [13] it was a new observation for the R466A/K467A mutant; reinforcing the notion that PDK1 activation followed closely its homodimerisation status.

Taken together, these data suggested that growth factor stimulation was promoting the formation of active PDK1 homodimers through binding to PtdIns(3,4,5)P_3_ at the plasma membrane. We suggest an alternative mechanism whereby the formation of active PDK1 homodimers may bypass binding to PtdIns(3,4,5)P_3_. We propose that PtdIns(3,4,5)P_3_ and other anionic phospholipids concurred to localise PDK1 at the PM but their differential binding triggered opposite effects on PDK1 regulation. PtdIns(3,4,5)P_3_-binding induced a conformation that was conducive to PDK1 and downstream substrates activation, whereas the binding to anionic phospholipids other than PtdIns(3,4,5)P_3_ maintained PDK1 in an inactive conformation at the PM.

To understand the dynamics of anionic phospholipids associated to the mechanism of PDK1 regulation, we took advantage of a cell line with a dysregulated PI3 kinase and an altered phosphoinositide composition. SKBR3 cells are a breast cancer cell line with over expressed levels of HER2 inducing anomalies in the PI3 kinase pathway. First we investigated the modulation of PtdIns(3,4,5)P_3_ by growth factor stimulation and PI3K inhibition and the PtdSer levels in these cells. This in turn was compared to how these anionic phospholipids affected the formation of dimers of PDK1 and its mutants in SKBR3.

### Determination of the relative *in situ* levels of PtdIns(3,4,5)P_3_ by GRP1 PH domain

To test how the aberrant behaviour of the HER2 growth factor receptor pathway, in SKBR3 cells, affected the levels of PtdIns(3,4,5)P_3_, we used an *in situ* intensity segmentation method to quantify PtdIns(3,4,5)P_3_ levels. In these experiments, we specifically chose a PH domain probe with a relatively low affinity for PtdIns(3,4,5)P_3_ detection (GFP-GRP1^PH^ [27]). The lower affinity of this probe (*K*_D_≈50 nM [28]), compared to PH^PDK1^, (*K*_D_≈0.2 nM [29]) reduced the cytoplasmic sequestration of PtdIns(3,4,5)P_3_ and thus provided a more accurate representation of endogenous levels of PtdIns(3,4,5)P_3_ at the plasma membrane. Compared to NIH3T3 cells, PtdIns(3,4,5)P_3_ levels at the plasma membrane were higher, in SKBR3 cells, upon stimulation of the PI3K pathway (Fig. S5). The PI3K inhibition (LY294002) prior to stimulation resulted in the decrease of PtdIns(3,4,5)P_3_ levels at the plasma membrane below the basal levels observed in NIH3T3 cells. PtdIns(3,4,5)P_3_ levels remained at the basal level in SKBR3 cells. These data suggested either an additional PI3K independent source of PtdIns(3,4,5)P_3_ or, that elevated PtdIns(3,4,5)P_3_ was a consequence of enhanced receptor tyrosine kinase activity in SKBR3 cells and the residual uninhibited PI3K. The control binding-defective GRP1 PH domain mutants did not bind to PtdIns(3,4,5)P_3_.

These results were also independently confirmed in live cell experiments using the same quantification method for PtdIns(3,4,5)P_3_. That is, PtdIns(3,4,5)P_3_ levels increased upon stimulation at the plasma membrane and were reduced below basal level upon PI3K inhibition in NIH3T3 but not in SKBR3 cells (Fig. S6). The GFP-GRP1^PH^ probe followed the same behaviour in starved and non-starved SKBR3 cells (Supporting Fig. S7).

PtdSer levels in both cell lines were also quantified using a genetically encoded fluorescently labelled PtdSer sensor (mCherry-Lact-C2) [30] fused to a self-cleaving P2A peptide and to GFP [31]. This way mCherry-Lact-C2 and free-diffusing GFP were expressed at the same rate, allowing ratiometric imaging of the sensor. We did not observe any growth factor stimulation-related differences in localisation to the plasma membrane of the PtdSer biosensor (Fig. S7). In summary, these data demonstrated that the levels of PtdSer were not significantly modified by stimulation or inhibition of the PI3K pathway in both cell lines. The regulation of the PtdIns(3,4,5)P_3_ levels was altered in SKBR3 cells. Unlike NIH3T3 cells, the basal level of PtdIns(3,4,5)P_3_ in SKBR3 was not affected by PI3K inhibitor (LY294002).

### In dysregulated PI3K cells, PDK1 homodimerisation and localisation was independent of its PH binding to PtdIns(3,4,5)P_3_ and PtdIns(3,4,5)P_3_ levels

Having measured the changes in PtdIns(3,4,5)P_3_ levels between the two cell lines, we next monitored the dynamic regulations of PDK1 in SKBR3 cells, where a different behaviour from NIH3T3 cells was identified. All PDK1 variants showed a strong constitutive association with the plasma membrane (Fig. 3A and B); this was also consistent for the PtdIns(3,4,5)P_3_ binding-defective (K465A) and the R466A/K467A PDK1 mutants. The PI3K inhibition prior to stimulation did not affect the recruitment of any of the PDK1 variants to the SKBR3 plasma membrane (Fig. 3B). A prominent PM localisation was observed for the homodimer conformers, including mutant R466A/K467A-PDK1. In the NIH3T3 cells, these were localised preferentially in the cytoplasm (Fig. 3D). Our results indicated that in NIH3T3 cells the localisation of PDK1 (irrespective of its dimerisation status) was dependent on the transient production of PtdIns(3,4,5)P_3_ at the PM. But in SKBR3 cells, PDK1 localisation was constitutively at the PM and did not change significantly, either through PDK1 PH domain mutation (K465A) or upon PI3K stimulation or inhibition. This suggested that the overall localisation of PDK1 was independent of the transient formation of PtdIns(3,4,5)P_3_. Furthermore, the double mutation R466/K467 to alanine did not prevent PDK1 recruitment to the PM, indicating events other than anionic lipids binding to the PH domain were involved in PDK1 recruitment.

**Fig.3.**
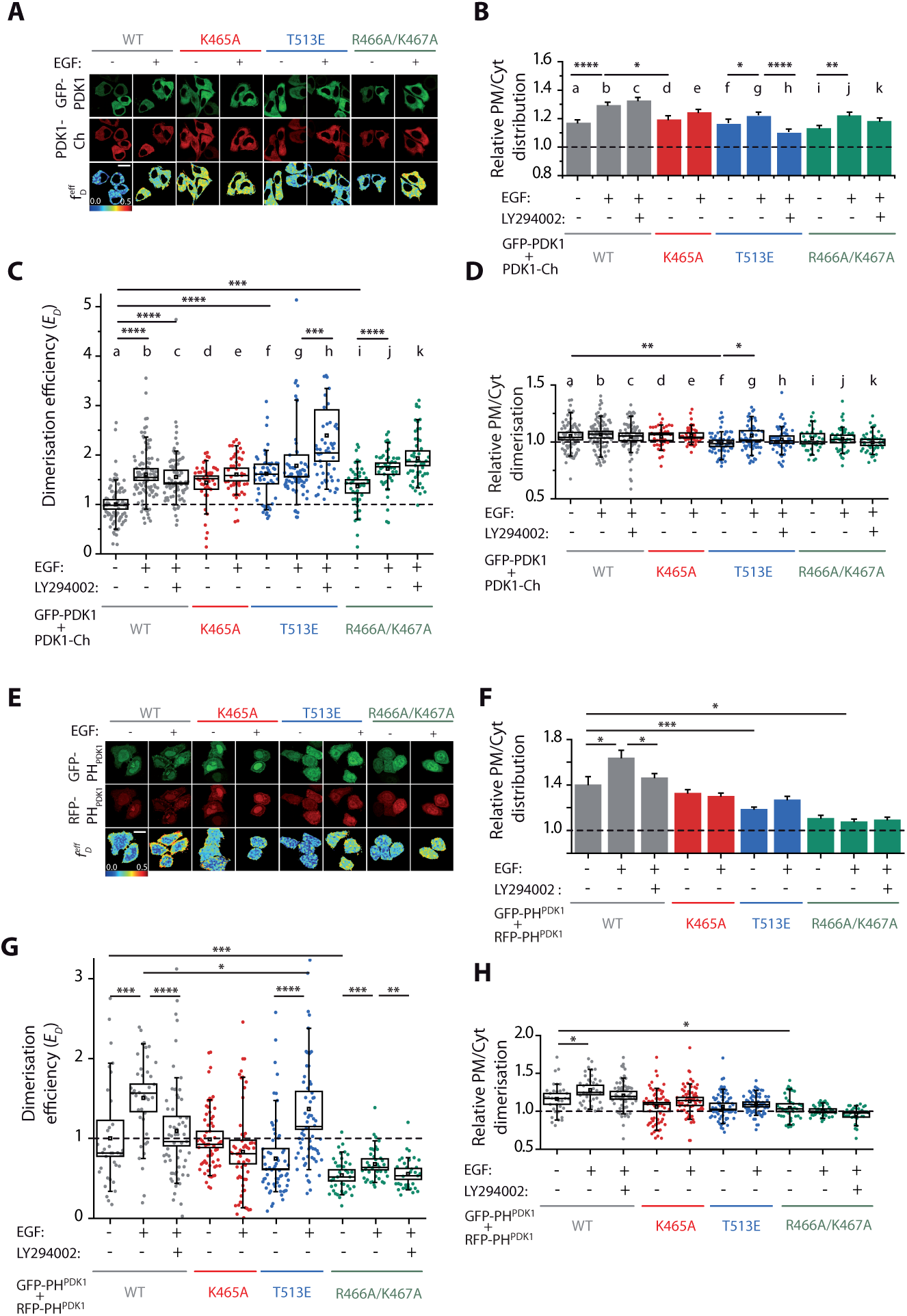
In dysregulated PI3K cells, PDK1 homodimerisation and localisation is independent of its PH domain binding to PtdIns(3,4,5)P_3_ and PtdIns(3,4,5)P_3_ levels. **(A-D)** SKBR3 cells were co-transfected with GFP-PDK1 and PDK1-mCherry **(E-H)** SKBR3 cells were co-transfected with GFP- and mRFP-tagged PH^PDK1^, in resting (-) and growth-factor (+) stimulated conditions. **(B and F)** Translocation of full-length PDK1 (B) or PH^PDK1^ (F) to the PM is quantified as the average fluorescence signal per pixel at the PM relative to the cytoplasm. **(C and G)** Homodimerisation efficiency of ectopic WT and mutants full-length PDK1 (C) or PH^PDK1^ (G) upon stimulation and PI3K inhibition. **(D and H)** The localisation of full-length PDK1 (D) or PH^PDK1^ dimers (H) was determined by the segmentation of the FRET images and is represented by the relative PM/cytoplasm distribution. Scale is 20 µm. N>30. Error bars: SEM. (C) and (D) N>30. Box: 2xSEM (95.4% confidence); Whiskers: 80% population. Mann-Whitney test *p<0.05. Three independent experiments.

Our next step entailed interrogating how the constitutive PM localisation would influence the formation of PDK1 homodimers in dysregulated SKBR3 cells. Fig 3C shows that WT-PDK1 responded to EGF stimulation shown by the increase of the homodimer population. However, PI3K inhibition did not prevent the EGF-stimulated increase in WT-PDK1 homodimers (Fig 3Cc). This result was not due to a lack of effect of the PI3K inhibitor since it blocked EGF-induced PtdIns(3,4,5)P_3_ increase in SKBR3 cells (Fig. S5h). It implied that while the increase in homodimers was dependent on EGF stimulation, it was independent of transient formation of PtdIns(3,4,5)P_3_. The unexpected behaviour of the PDK1 mutants showed a significant increase in the basal homodimerisation compared to WT-PDK1.

The fact that PDK1 was able to homodimerise independent of variations in PtdIns(3,4,5)P_3_, and that none of the PH domain mutations affected the binding of full-length PDK1 to the PM, pointed towards a critical role for the kinase domain in the regulation of PDK1 localisation and dynamics. To test this hypothesis, we investigated how isolated PDK1 PH domains in SKBR3 cells would recruit and dimerise at the plasma membrane. The cells were co-transfected with GFP-PH^PDK1^ and mRFP-PH^PDK1^ (Fig. 3E). The isolated PH domains of PDK1 localised at the PM analogous to full-length PDK1 in these cells. Contrary to the full-length protein their relative PM/Cytoplasm distribution was modulated by PtdIns(3,4,5)P_3_ varying levels upon growth factor stimulation and PI3K inhibition. The (K465A) PH-domain mutant responded accordingly. All the mutations affected the localisation of the PH domains to the PM, with the mutation R466A/K467A having the strongest effect.

Direct assessment of the wild type PH^PDK1^ homodimerisation in SKBR3 cells showed that it was PtdIns(3,4,5)P_3_-dependent, contrary to the full-length protein (Fig. 3G). EGF stimulation induced the production of WT-PH^PDK1^ dimers, inhibited by pretreatment with LY294002. The loss of PtdIns(3,4,5)P_3_ binding in the mutant (K465A) prevented the formation of homodimers upon stimulation. Similar to the wild type, the activating mutant PH domain (T513E) dimers were formed upon stimulation. The mutation of the R466/K467 residues totally abolished the basal dimerisation of the PDK1 PH domains. The PH domain homodimer population (Fig. 3H) was localised at the plasma membrane and was reduced by LY294002 treatment or mutation of the PH domain. The loss of the homodimers at plasma membrane was most significant in the case of the R466A/K467A-PH^PDK1^ mutant.

Altogether these data revealed that the dynamic regulation of the isolated PH domains of PDK1 was different from the regulation of full-length PDK1 in SKBR3 cells. PH^PDK1^ variants homodimerised to a level that was consistent with their localisation at the plasma membrane and the transient production of PtdIns(3,4,5)P_3_ (Fig. 3E-H). The full-length protein did not behave in this manner Fig. 3C). Therefore, these results indicate that the PH domain was not the sole domain with a role in the localisation of full-length PDK1 at the plasma membrane or its homodimerisation. We highlight the critical contribution of the kinase domain in the modulation of PDK1 conformational dynamics. Furthermore, the differences between the full-length PDK1 and isolated PH domain illustrated that in SKBR3 cells, PDK1 was not exclusively regulated by the acute formation of PtdIns(3,4,5)P_3_. Instead, other anionic phospholipids would concur to the regulation of PDK1 in a kinase domain-dependent manner.

Our data demonstrates that PDK1 dimerisation by the modulation of PtdIns(3,4,5)P_3_ levels was the major determinant of PDK1 activity in NIH3T3 cells (Fig. 2D). In SKBR3 cells where PI3K pathway is dysregulated, the formation of PDK1 dimers (Fig. 2G, c) did not concur with PtdIns(3,4,5)P_3_ variations (Fig. 3C, c, h, k). We suggest that the conventional model for the regulation of PDK1 activity by the transient production of PtdIns(3,4,5)P_3_ is incomplete, as it does not account for the regulation of PDK1 in cells with a dysregulated PI3 kinase pathway.

### PDK1-Akt/PKB heterodimerisation and downstream activation of Akt/PKB correlates with PDK1 homodimer formation in SKBR3 cells

To determine how the aberrant production of PDK1 homodimers in SKBR3 cells influenced downstream substrates activation, we monitored the phosphorylation of two PDK1 substrates Akt/PKB and SGK1. Previously, we had shown that Akt/PKB activation required PDK1-Akt/PKB heterodimerisation after binding to PtdIns(3,4,5)P_3_ [10]. We proceeded in determining their interaction by coexpressing, in SKBR3 cells, eGFP-myc-PDK1 and mCherry-Akt/PKB (Fig. 4A and Fig. S1C). We determined that the population of PDK1-Akt/PKB heterodimers (Fig. 4B) correlated with the formation of PDK1 homodimers observed in Figure 3C. Similar to PDK1 homodimerisation, the heterodimer population of WT-PDK1-Akt/PKB increased upon EGF stimulation. The PtdIns(3,4,5)P_3_ binding-defective mutation K465A did not prevent the heterodimerisation of PDK1 with Akt/PKB. Following the same pattern than WT-PDK1, the interaction of T513E-PDK1 with Akt/PKB showed an enhanced heterodimerisation upon activation. We demonstrated the significance of the R466A/K467A mutation in signalling propagation, the binding of the R466A/K467A-PDK1 mutant to Akt/PKB was significant (Fig. 4B g, h), which again correlated with the elevated *E*_D_ of the PDK1 homodimer detected for the same mutant (Fig. 3C).

**Fig.4.**
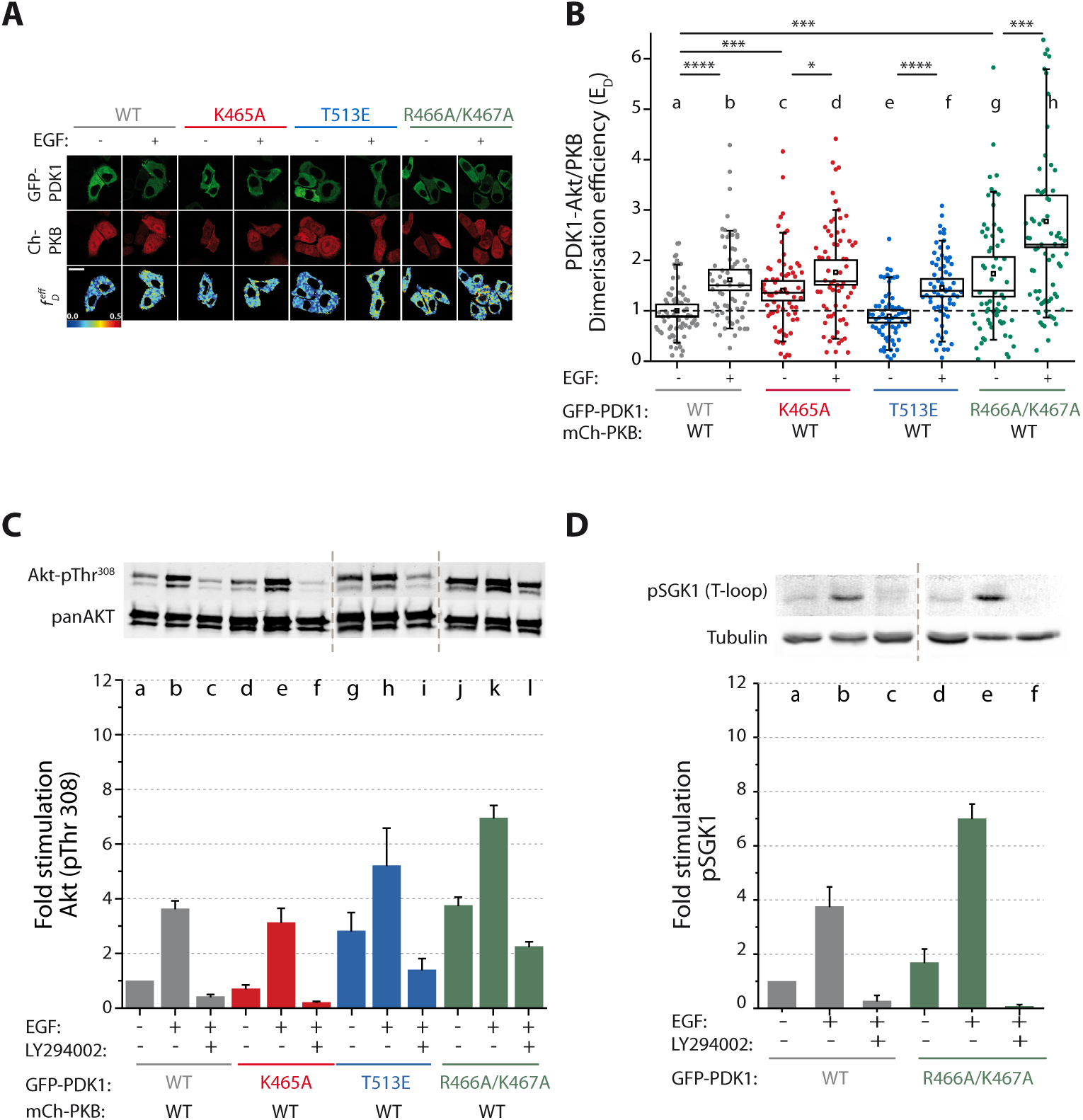
Akt/PKB heterodimerisation with PDK1 and downstream activation correlates with the level of PDK1 homodimerisation in SKBR3 cells. **(A)** Representative intensity and FRET-FLIM images of SKBR3 cells co-transfected with mCherry-Akt/PKB and GFP-PDK1. **(B)** The efficiency of mCherry-Akt/PKB and GFP-PDK1 interaction determined by FRET-FLIM in SKBR3 cells increases upon growth-factor stimulation and is highest for R466A/K467A PDK1 mutant. **(C)** Phosphorylation of Akt/PKB at T308 correlates well with the efficiency of Akt/PKB and PDK1 heterodimerisation. **(D)** Phosphorylation of endogenous SGK1 at its T-loop by PDK1 is potentiated by the mutation R466A/K467A. (A) Scale bar: 20 µm. (B) N>30. Box: 2xSEM (95.4% confidence); Whiskers: 80% population. Mann-Whitney test *p<0.05. Three independent experiments. (C), (D) and (E) n=3. Error bars: SEM

The enhanced interaction of all the PDK1 variants with Akt/PKB upon stimulation (Fig. 4B, b, c, f, h) led in each case to an increase of Akt/PKB phosphorylation at its activation loop T308 (Fig. 4C). PI3K inhibition induced a reduction in PtdIns(3,4,5)P_3_ levels in SKBR3 (GRP1 probe - Fig. S5) and also inhibited Akt/PKB T308 phosphorylation upon stimulation. The PDK1 mutation K465A, did not prevent the EGF-triggered increase in T308 phosphorylation. This was consistent with the fact that this mutation did not obstruct either the localisation of full-length PDK1 K465A to the plasma membrane (Fig. 3C) or the increase in PDK1-Akt/PKB heterodimerisation (Fig. 4B). A significant increase in T308 phosphorylation was observed with both the activating mutant T513E and the R466A/K467A mutant compared to WT-PDK1, specifically under basal conditions (Fig. 4C, g, j). This correlated with the fact that T513E increased PDK1 activity and that the R466A/K467A mutation induced the formation of PDK1-Akt/PKB heterodimers in basal conditions (Fig. 4B).

In order to test whether the activation of PDK1 due to the mutation of R466A/K467A could also impact more endogenous substrates other than Akt/PKB, we investigated the phosphorylation status of endogenous SGK1 at its T-loop (T256) (Fig. 4D and Fig. S8). Activation of SGK1 is known to be dependent on PI3K activation and the production of PtdIns(3,4,5)P_3_ and requires its hydrophobic motif phosphorylation by mTORC2 [32, 33]. However, unlike Akt/PKB, the kinase itself does not have a PH domain to directly interact with PtdIns(3,4,5)P_3_ [33]. Figure 4D shows a similar effect seen for Akt/PKB. We observed the enhanced phosphorylation of SGK1’s T-loop upon stimulation with EGF in a PI3K-dependent manner upon expression of WT PDK1. Furthermore, as observed with AKT/PKB, PDK1 R466A/K467A mutant also increased the phosphorylation of SGK1 under basal and stimulated conditions (Fig. 4D).

Altogether, the data obtained in both NIH3T3 and SKBR3, suggest that in addition to the well-established requirement for PtdIns(3,4,5)P_3_, the binding of PDK1 to other anionic phospholipids was also critical in the regulation of PDK1 downstream substrates. More specifically, while PDK1 interaction with PtdIns(3,4,5)P_3_ would trigger downstream substrate activation, the binding to anionic phospholipids through K466/R467 would be inhibitory.

### Molecular conformations of PDK1 PH domains homodimers binding to PtdIns(3,4,5)P_3_ and/or PtdSer

Molecular modelling and docking experiments were performed (Supplementary Information) to understand the molecular mechanisms whereby PDK1 dynamics would be differentially regulated by binding to PtdIns(3,4,5)P_3_ and PtdSer. This approach also determined the binding site for these anionic phospholipids. The binding of PtdIns(3,4,5)P_3_ and PtdSer were modelled on isolated PDK1 PH domain. The experiments showed that both anionic phospholipids recognised the same site on the PDK1 PH domain, albeit through different interactions and affinities (Fig. 5A). These results indicated that contrary to previous propositions [6] the two phospholipids could not bind concomitantly to the same PH domain. The obstruction of PtdSer in the binding of PDK1 PH domains in GUV containing PtdIns(3,4,5)P_3_ (Fig.1), was therefore unlikely to be due to the two phospholipids binding to the same PH domain. An alternative explanation was that PH^PDK1^ could bind to PtdSer and PtdIns(3,4,5)P_3_ as a dimer with each PH monomer binding to a different phospholipid. By performing protein-protein docking experiments we identified three possible PH domain homodimer conformers (C1, C2 and C3) formed by interacting with different combination of PtdSer and PtdIns(3,4,5)P_3_ binding to the PH domain pocket (Fig. 5B). The unstructured N-terminal linkers that mediate association to the kinase domains are also presented in Figure 5B to evaluate the positioning of the kinase domain in the various conformers (red segments). The C1 conformation is associated only with one PtdSer molecule bound to one of the PH domains in the dimer while the other PH domain remained unoccupied. In C1, the relative positioning of the two PH domains did not allow for the simultaneous binding of two phospholipids at the same time at the membrane. Furthermore, the localisation of the linkers to the kinase domains suggested a close proximity of the two kinases (see the closed angle of 49°). Similar to C1, the C3 conformer arises when only one lipid was bound to one PH domain in the dimer, but in the case of C3 the lipid was PtdIns(3,4,5)P_3_. Both C1 and C3 could potentially adopt a C2 conformation by binding to either another PtdSer or a PtdIns(3,4,5)P_3_ with the empty remaining PH domain. The C2 conformation was compatible with a full access to both lipid sites at the membrane. Contrary to the C1 conformation, the linkers in C2 were at maximum distance and angle (178°). This shows that the kinase domains in C2 were much further apart than in the C1 conformation, and could be in a position allowing the access of the substrates to the kinase active sites. In this condition, the C2 conformer would be compatible with an active conformation of the PDK1 homodimer. It is worth noting that while the angles between the conformers C2 and C3 are the same, the complexes bear a different geometry and might not have the same activity.

**Fig.5.**
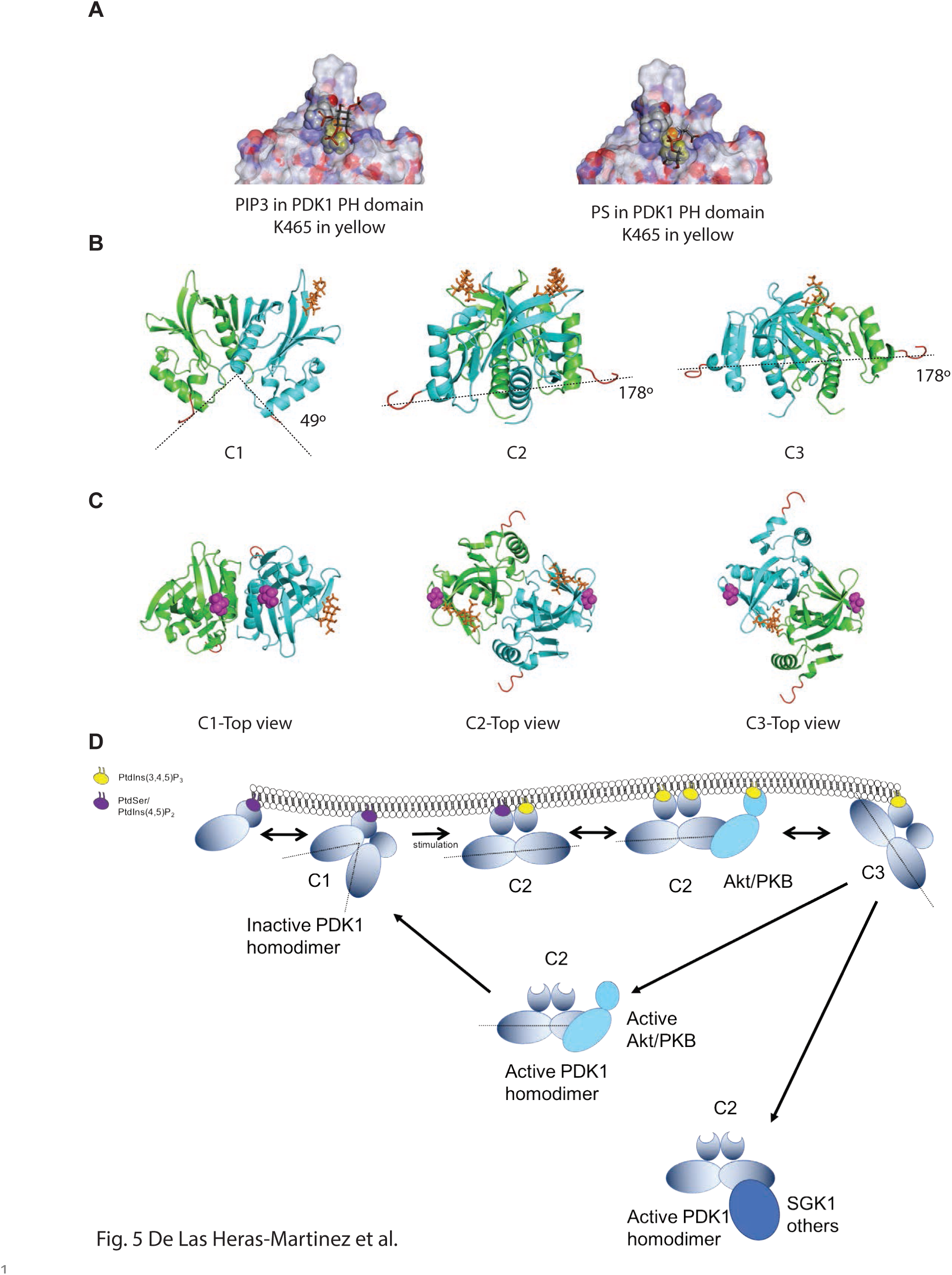
Molecular modelling of PDK1 PH domain homodimers and model of PDK1 homodimers activation. **(A)** Docking of PtdIns(3,4,5)P_3_ (left panel) and PtdSer (right panel) on PDK1 PH domain. Residue K465 is in yellow. **(B)** Representation of the three possible PDK1 PH domain homodimers bound to anionic phospholipids. **(C)** Model of the predicted active and inactive PDK1 homodimer conformers and their interaction with downstream substrates.

In light of these results, we suggest (Fig. 5C) that prior to stimulation, PDK1 homodimers would bind to the abundantly available PtdSer (and from our biochemical data possibly also PtdIns(4,5)P_2_) in an inactive C1 conformation. This event would permit PDK1 recruitment to the plasma membrane, but due to close proximity of the two kinase domains, C1 conformation would remain inactive. Upon growth factor stimulation and increase in PtdIns(3,4,5)P_3_ levels at the plasma membrane, the further interaction of C1 with PtdIns(3,4,5)P_3_ via its unoccupied PH domain would trigger a change in conformation towards an active C2 conformer. The separation of the two kinases in the C2 homodimer would allow accessibility to substrates and hence downstream activation. The molecular modelling data also predicted that both the binding of two PtdIns(3,4,5)P_3_ or a non-lipid-binding state (where both PH domains are empty) would also trigger the active C2 conformation of the PDK1 homodimers. These results were of great significance since they were compatible with our biochemical findings where both PtdIns(3,4,5)P_3_-triggered active plasma membrane homodimers as well as cytoplasmic (unbound) PDK1 homodimers, were capable of phosphorylating downstream substrates like Akt/PKB and SGK1.

The molecular mechanisms that would lead to the disassociation from the plasma membrane have not yet been identified. Our work and others suggest that it would likely involve the autophosphorylation of T513 and S410. We suggest that electrostatic repulsion would be a likely mechanism for triggering the loss of electrostatic binding to anionic phospholipids. Our molecular modelling data (Table 1) showed that the phosphorylation of T513 prevented the formation of the C1 conformer, suggesting that the presence of phospho-T513 would not be compatible with an inactive conformation of the PDK1 homodimer. Table 1 also shows that in the C3 conformer both S410 and T513 are closer to the inositol binding site than other conformations. This suggests that C3 might be a transition conformer that would occur following the loss of binding to one of the PtdIns(3,4,5)P_3_. The autophosphorylation of the residues S410 and T513 in the C3 (positioned closer to the anionic phospholipid binding site than in other conformations) could trigger repulsion from the plasma membrane.

**Table 1:** Distances between residues approximated to the closest number.

| Monomer A-Monomer B<br>distances (angstroms) | C1 | C2 | C3 |
| --- | --- | --- | --- |
| Ser410 – Binding pocket | 44 | 33 | 31 |
| Ser410 – Thr513 | 40 | 39 | 41 |
| Thr513 – Binding Pocket | 30 | 37 | 29 |
| Thr513 – Thr513 | 15 | 48 | 43 |
| Ser410 – Ser410 | 31 | 52 | 64 |

## Discussion

Previously, we identified PDK1 homodimerisation as a central component of its acute regulation [13]. We showed by *in situ* FRET-FLIM that PDK1 homodimerised in the cytoplasm of COS7 cells in a PtdIns(3,4,5)P_3_ dependent manner. Furthermore, we suggested that the autophosphorylation of T513 upon stimulation might lead to the production of active PDK1 monomers through disruption of the inactive homodimers [13].

The current refinement of our FRET-FLIM methodology [23] permitted a deeper understanding of the complex mechanisms underpinning the regulation of PDK1. By monitoring the conformational dynamics of the PDK1 T513E mutant *in situ* in NIH3T3 (Fig. 2B) and SKBR3 cells (Fig. 3C) we observed that the T513E mutation did not lead to the disassembly of the PDK1 homodimers, but generated an increase in a homodimer population correlating with the activation of one of its downstream substrates (Akt/PKB) (Fig. 2D and Fig. 4B, C). This may be interpreted in two ways. Firstly, that the PDK1 conformer requires a transition to an activated homodimer conformation prior to downstream substrate phosphorylation. Secondly, that PDK1 homodimerises in more than one conformation. We suggest that PDK1’s autophosphorylation at the T513 residue, triggered upon plasma membrane translocation, could be a switch or a stabilising factor of the conformational change. Work by Kang *et al.* suggested, using co-immunoprecipitation of the isolated PDK1 PH and kinase domains, the presence of more than one possible PDK1 homodimer conformation [22]. They proposed a key role for the residue T513 in the formation of different dimers.

Our recent quantification of the PDK1 homodimerisation affinity determined by the effective intracellular dissociation constant 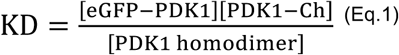 in SKBR3 cells, also demonstrated the presence of more than one homodimer conformation [23]. In this latter work, PDK1 homodimerisation was detected by time resolved-FRET between GFP- and Cherry-tagged PDK1 and the total concentration of PDK1-Cherry and GFP-PDK1 was determined by intensity measurements. The FRET measurements determined the concentration of PDK1 homodimers. From these measurements and using (Eq.1), we established that the effective intracellular dissociation constants at basal conditions ((K_D_)_basal eff_ = 49 μM) were greater than post stimulation ((K_D_)_EGF eff_ = 11 μM). The explanation for the decrease in the effective K_D_ upon stimulation could be the contribution of a “FRET-inactive” PDK1 homodimer population (a dimer population that does not FRET), versus the FRET-active PDK1 homodimer population. The “FRET-inactive” dimer would correspond to a conformer geometry unfavourable to the efficient transfer of energy. This, can issue from the variability in the orientation of the chromophores, a parameter which in addition to the distance also defines efficient transfer of energy. The results from the de Las Heras paper [23] supports the idea of PDK1 homodimer species with different conformations, coexisting at equilibrium with their relative abundance being dependant on the activation signal.

We suggest that the homodimer conformers with different topologies are a more effective mechanism of regulating PDK1 rather than the pre-assumed monomeric forms. This effective mechanism may provide a refined control of PDK1 activity by permitting a broader array of anionic phospholipids binding to the various topologies. The occurrence of inactive PDK1 homodimers could be explained by a transition ‘primed’ conformer species. In this conformation, the PDK1 conformer would be phosphorylated on the activation loop but auto-inhibited prior to stimulation. This would allow for a rapid activation of PDK1 upon growth factor stimulation. Such behaviour is well established in other AGC kinases, such as PKCs. The members of this family are fully phosphorylated ‘primed’ at their activation sites but autoinhibited by a pseudosubstrate sequence prior to stimulation [34]. PDK1 has been shown to be constitutively autophosphorylated on its activation loop (S241) independently of growth factor stimulation and therefore phosphorylated on this residue in its inactive as well as active conformations [18]. Our molecular model (Fig. 5) provides the possible conformation of an inactive PDK1 homodimer, prior to PtdIns(3,4,5)P_3_ binding and autoinhibitory configuration where the close proximity of the kinases would hinder the interaction with substrates (Fig. 5 dimer C1). The possibility of dimeric PH domains allows a combinatory binding of various anionic lipids that would not be feasible with monomeric PDK1, thus providing a refined regulatory step for its activation.

By exploiting the spatial and conformational dynamics of PDK1 and its PH domain mutants (K465A and R466A/K467A), we identified the differential role of the binding of anionic phospholipids as a molecular switch of the inactive to the active PDK1 conformations. The *in vitro* affinity measurements of isolated PDK1 PH domains indicated that the affinity for PtdSer is two to three orders of magnitude lower than for PtdIns(3,4,5)P_3_ (Fig.1 A versus B; Fig. 1C). We showed that the mutation K465A inhibits the binding to PtdIns(3,4,5)P_3_ and PtdIns(4,5)P_2_ but not PtdSer, and that conversely the mutation R466A/K467A prevents the interaction of PtdSer and PtdIns(4,5)P_2_ but not PtdIns(3,4,5)P_3_. Therefore, despite the close proximity of the residues K465, R466 and K467 in PDK1 PH domain, the role of different anionic phospholipids in cells can be distinguished by using these differential mutations. The results from the FCS experiments using GUVs, confirm the specific binding of PDK1 PH domain to PtdIns(3,4,5)P_3_ containing GUVs. The presence of PtdSer in the same GUV interfered with the PH^PDK1^ binding to PtdIns(3,4,5)P_3_, suggesting that the close proximity of PtdSer to the PtdIns(3,4,5)P_3_–bound PH^PDK1^ facilitated the association of PtdSer. Since our molecular model (Fig. 5A) established that the docking cavity cannot simultaneously accommodate PtdIns(3,4,5)P_3_ and PtdSer as previously proposed [6], our results suggest that PDK1 may bind both lipids as a dimer, with each PH unit binding a different lipid.

Our model demonstrates that PtdIns(3,4,5)P_3_ and PtdSer bind to a PDK1 PH-PH dimer, with each anionic phospholipid binding to one PH domain. Moreover, it predicts the formation of three types of PH^PDK1^ homodimers with various propensities to interact with anionic phospholipids. Briefly, the C1 conformer binds to PtdSer (only one of the two PH binds); C2 conformer binds to two anionic phospholipids, (PtdSer-PtdIns(3,4,5)P_3_) or (PtdIns(3,4,5)P_3_-PtdIns(3,4,5)P_3_); the C3 conformer binds only to PtdIns(3,4,5)P_3_ (only one of the two PH binds).

In addition to the regulation by anionic phospholipids, the post-translational modification of the PH domains by T513 phosphorylation can affect PDK1 activation in a PH domain dependent manner. Binding affinity assays and FCS data showed that the activating PH domain mutation T513E induced the loss of PH^PDK1^ specific binding to PtdIns(4,5)P_2_ and PtdSer and a strong reduction to PtdIns(3,4,5)P_3_ (Fig. 1B-D). Therefore, suggesting the activating effect of T513 phosphorylation would be through the disassociation of PDK1 from the plasma membrane, thereafter promoting PDK1 interaction with its downstream substrates.

By monitoring the localisation and the conformational dynamics of full-length PDK1 in NIH3T3 cells, by FRET-FLIM, we confirmed the interplay of the anionic phospholipids. PDK1 plasma membrane translocation upon stimulation induced a PtdIns(3,4,5)P_3_ binding-dependent increase in homodimerisation (mutant K465A). This increment was correlated with an increase in downstream substrate phosphorylation. These results suggested that activated PDK1 homodimers, upon stimulation, were involved in the activation of the downstream substrates. We observed that the loss of PM binding (R446A/K467A mutant), also elicits a strong constitutive dimerisation of PDK1 promoting downstream substrates phosphorylation (Fig. 2 B, D). These data implied that PDK1 homodimer formation occurs through two independent mechanisms; one dependent on the binding to PtdIns(3,4,5)P_3_ at plasma membrane and the other in response to the loss of the plasma membrane binding (through the R446A/K467A mutation). In both cases, the formation of homodimer is correlated with the activation of its downstream substrate Akt/PKB. This suggested that the homodimers are similar in conformation and their activation state. We propose that the binding to PtdIns(3,4,5)P_3_ is responsible for the activation of PDK1 by forming PDK1 activated homodimers. Conversely, the anionic phospholipids binding to residues R466/K467 prior to PtdIns(3,4,5)P_3_ formation (in basal conditions), are therefore responsible for maintaining PDK1 in an inactive conformation.

The abundance of PtdSer and PtdIns(4,5)P_2_ and the fact that PtdSer can cooperate with PtdIns(3,4,5)P_3_ to target PDK1 PH domain to the plasma membrane (Fig 1 D), as observed in other PH domains [35, 36], suggests that these anionic phospholipids are responsible for the negative regulation of PDK1 through binding to R466/K467. Furthermore, the R466A/K467A mutant resulted in a loss of its specific cellular localisation without a significant effect on the formation of total dimer population. This implies that the homodimers of PDK1 are differentially regulated by anionic phospholipids, a prerequisite for the refined regulation of PDK1. The association with PtdSer (and possibly PtdIns(4,5)P_2_) increases the membrane “residence time” of the full-length protein and may affect its affinity for PtdIns(3,4,5)P_3_. These effects could provide PDK1 with a search mechanism for PtdIns(3,4,5)P_3_, similar to that proposed for GRP1^PH^ [37]. This regulation mechanism would also be compatible with the micro-domain compartmentalisation during the initial stages of PI3K signalling and the constrained lateral diffusion exhibited by PDK1’s immediate downstream target Akt/PKB [38, 39].

To determine the effect of the dysregulation of the PI3K pathway on the regulation of PDK1, we utilised SKBR3 cells that have constitutively elevated PtdIns(3,4,5)P_3_ levels. We showed that PDK1 regulation was perturbed in this cell line. Growth factor stimulation still resulted in activated PDK1 homodimer formation (Fig. 3C). Binding and activation of the downstream effectors Akt/PKB and SGK1 (Fig. 4C, D) was also triggered. However, the inhibition of PI3K pathway due to the pre-elevated PtdIns(3,4,5)P_3_ levels did not affect dimer formation, suggesting that under dysregulated conditions, PtdIns(3,4,5)P_3_ does not behave as the molecular switch. Essentially, all mutants K465A, R466A/K467A and T513A elicited a PDK1 homodimerisation, which correlated with downstream substrates activation, indicating the formation of activated homodimers. The overall population of PDK1 homodimers was mainly located at the plasma membrane, irrespective of the PH domain mutations. This suggested that the kinase domain had a significant influence on the localisation and dynamics of a cellular environment exhibiting a dysregulated PI3K pathway (SKBR3 cells). Contrary to the full-length PDK1, the PH^PDK1^ wild type and its mutants behaved as predicted in a regulated PI3K pathway (as in NIH3T3 cells). Therefore, the kinase domain is a major determinant in the regulation of PDK1 not only in terms of conformational dynamics but also through its localisation at the plasma membrane.

This finding corroborates with Komander *et al.* who have also suggested that full-length PDK1 localisation in cells did not strictly follow the occurrence of PtdIns(3,4,5)P_3_ at the plasma membrane unlike the isolated PDK1 PH domains [5]. They postulated that PDK1 could be sequestered in the cytosol possibly by binding to soluble inositol phospholipids IP5/IP6. It is not clear whether this sequestration would involve the binding of monomers or dimers of PDK1, and how the presence of the kinase domain would influence this sequestration but it could be part of the mechanism that distinguishes the behaviour of PDK1 in SKBR3 from NIH3T3 cells.

In summary, our results reveal that PDK1 homodimerised in response to an increase in PtdIns(3,4,5)P_3_ levels and that the activated PDK1 dimers phosphorylated both membranous (Akt/PKB) and cytoplasmic (SGK1) substrates. Our observations establish the cooperative association of PtdSer (and possibly other anionic lipids like PtdIns(4,5)P_2_) with PtdIns(3,4,5)P_3_ in recruiting PDK1 to the plasma membrane but they have opposite effects on PDK1 activation. Our results indicate that membrane-binding to anionic lipids other than PtdIns(3,4,5)P_3,_ would stabilise plasma membrane-associated PDK1 in a homodimer conformation where the catalytic site would be sequestered from potential substrates. The enhancement in PtdIns(3,4,5)P_3_ levels would induce a dual binding of the PDK1 homodimer from PtdSer only (C1) to PtdSer-PtdIns(3,4,5)P_3_ or PtdIns(3,4,5)P_3_-PtdIns(3,4,5)P_3_ and a change in conformation of PDK1 consistent with an activated conformation (C2). The phosphorylation of T513 leads to a transition C3 conformation and dissociation from the membrane where the PDK1 dimer adopts the C2 type activated conformation.

41 missense substitutions were found in PDK1 linked to various types of cancers (Cosmic database). Correlated to our results, one of these residues R466 was found mutated 3 times; in brain (R466Q), breast (R466Q) and colon (R466W) [40]. Our findings indicate that a physiological alteration of the R466/K467 site in cells that would mimic the effect of the mutant 466A/467A, and alter the binding to PtdSer or PtdIns(4,5)P_2_, would lead to the constitutive activation of PDK1 and potentially to a significant effect in different types of cancer. Therefore, detailed characterisation of this site, and its previously unappreciated role in PDK1 down regulation, would enable defining a novel therapeutic strategy where a constitutively activated dimer of PDK1 could be rendered inactive by small molecules that would drive its conformation towards an inactive membrane-bound conformer.

## Materials and Methods

- Cloning
- Cell culture and transfection
- Western blots
- Quantitative IR Western blot
- Protein purification
- Protein lipid overlay assay
- Scanning Fluorescence Correlation Spectroscopy
- Confocal and time-resolved imaging
- Preparation of fixed samples for time-resolved FRET experiments
- Image segmentation and calculation of the plasma membrane to cytoplasm partition coefficient
- Quantification of accessible PtdIns(3,4,5)P_3_ and PtdSer
- Quantification of the Dimerisation Efficiency (*E*_D_)
- Segmentation of FRET-FLIM images
- Molecular Modelling

### Cloning

Both mutants K465A and R466A/K467A of human PDK1 were obtained by site-directed mutagenesis of the eGFP-myc-PDK1 and HA-PDK1-mCherry constructs [13] with the QuickChange mutagenesis kit (Agilent Technologies, Inc.). The oligos used for the mutagenesis were as follows: sense, 5’-ggcccagtggatgcgcggaagggtttatttgc-3’ and antisense, 5’-gcaaataaacccttccgcgcatccactgggcc-3’ for the K465A mutant and sense, 5’-ggcccagtggataaggcggcgggtttatttgcaag-3’ and antisense, 5’-cttgcaaataaacccgccgccttatccactgggcc-3’ for the R466A/K467A mutant. To obtain the constructs of the isolated PH domain of PDK1 (residues 404-556 at the C-terminus of PDK1), this domain was first amplified from the full-length eGFP-myc-PDK1 construct by PCR. The oligos were designed to introduce a myc tag sequence between the fluorescent protein and the PH domain (sense, 5’-ggaagatctgcaatggaacagaaactcatctctgaagaggatctgccccagaggtcaggc-3’ and antisense, 5’-ctagtctagatcactgcacagcggcgtccgggtggctctggtatcg-3’). The whole myc-PDK1 sequence was removed from the original plasmid by enzymatic digestion with BglII and XbaI. The amplified myc-PH segment containing sites for BglII and XbaI at 5’ and 3’, respectively, was inserted in the digested plasmid to finally obtain: pCMV-eGFP-myc-PH^PDK1^. The mRFP-myc-PH^PDK1^ was obtained by substitution of the eGFP by mRFP. To do so, we PCR amplified the mRFP sequence from the pCMV-mRFP-HA-PKB plasmid using two oligos that included EcoRI and BglII sites to remove the eGFP from the original plasmid by digestion with those enzymes: sense, 5’-ccggaattcatggcctcctccgaggacgtc-3’ and antisense, 5’-ggaagatctagctgcaccggtggagtg. The K465A and R466A/K467A mutants of the isolated PH domain constructs were generated by site-directed mutagenesis using the same oligos as for the full-length protein. The mCherry-HA-PKB construct was cloned by PCR amplification of the mCherry sequence from the pCMV-HA-PDK1-mCherry plasmid with the oligos: sense, 5’-ccggaattcatggtgagcaagggcg-3’ and antisense, 5’-ggaagatctcttgtacagctcgtccatgccgccg-3’. By enzymatic digestion with EcoRI and BglII of the original construct pCMV-mRFP-HA-PKB we removed the mRFP to insert the PCR product containing EcoRI-mCherry-BglII and finally obtain pCMV-mCherry-HA-PKB.

The constructs for the PH domain of GRP1 and its double mutant (K273A/R284A) were obtained by isolating the C-terminal part (residues 240-399) of the protein containing the PH domain (residues 263-380) from the original pTriEX6-versatile plasmids kindly provided by G. Chung (Cell Biophysics Lab, CRUK, London). The amplified C-terminus was inserted in a pCMV5 plasmid and fused to eGFP. The oligos for the PCR were as follows: sense, 5’-ggaagatctgaaagtatcaagaatgagc-3’ and antisense, 5’-ctagtctagattatttcttattggcaatcctccttttc-3’.

mCherry-6Gly-eGFP were obtained from mCherry and eGFP containing plasmids. Three PCR were required to obtain mCherry-6Gly-eGFP: PCR1: sense, 5′-ctagctagcatggtgagcaagggcgaggaggataac-3′ and antisense, 5′-gctcaccatgccgccgccgccgccgcccttgtacagctcgtc-3′; PCR2: sense, 5′-tgtacaagggcggcggcggcggcggcatggtgagcaaggg-3′ and antisense, 5′-aaatatgcggccgctttacttgtacagctcgtccatgcc-3′. The products obtained from the first and second PCR were used as templates for a last step using the sense PCR1 and the antisense PCR2 primers. The resulting construct contains NheI and NotI sites at 5′and 3′ respectively. The Cherry-6Gly-eGFP fragment was now removed from the parental vector by enzymatic digestion using those sites and it was finally inserted into a pCMV backbone.

mCherry-LactC2-P2A-eGFP was obtained following a similar procedure to mCherry-6Gly-eGFP using mCherry-LactC2 [30] and mCherry-P2A-eGFP constructs as the templates for the initial reactions PCR1 and PCR2, respectively. The oligos were: PCR1: sense, 5′-tagctagcatggtgagcaagggcgaggaggataac-3′ and antisense, 5′-cagcaggctgaagttagtagcacagcccagcagctcc-3′; PCR2: sense, 5′-ggagctgctgggctgtgctactaacttcagcctgctg-3′ and antisense, 5′-cccaagcttttacttgtacagctcgtccatgccgagagtgatc-3′. For the third PCR reaction, the sense PCR1 and antisense PCR2 oligos were used and the final template was digested with NheI and HindIII and inserted in a pCMV backbone.

### Cell culture and transfection

SKBR3 and NIH3T3 cells were obtained from ATCC. SKBR3 and NIH3T3 cells were maintained in DMEM (Dulbecco’s modified Eagle′s medium) with GlutaMAX containing 1% penicillin/streptomycin (P/S) and 10% foetal bovine serum (FBS) or donor calf serum (DCS), respectively, at 37°C and 5% CO_2_ for SKBR3 or 10% for NIH3T3. For experiments, cells were seeded at 300,000 cells per well of a six-well plate (for Western Blotting) or per MatTek dish (for FRET/FLIM experiments).

Cells were co-transfected with 0.5 μg of donor DNA and 1 μg of acceptor DNA using Lipofectamine LTX & Plus Reagent in OptiMem containing GlutaMAX, following the protocol specified by the manufacturer. SKBR3 cells were incubated in this mixture for 4 h at 37°C and 5% CO_2_ while NIH3T3 were incubated for 3 h at 37°C and 10% CO_2_. The medium was removed and replaced by DMEM (with FBS or DCS and P/S) for another 3-4 h to let the cells recover. Finally, to serum starve SKBR3 cells the medium was replaced with DMEM containing only 0.2% of BSA and 1% P/S for at least 20 h. NIH3T3 cells were not serum starved. Experiments were performed about 24 h after transfection.

### Western blots

Cells were seeded at 300,000 per well of a 6-well plate, transfected, serum-starved and stimulated as indicated above. After stimulation cells were washed twice in cold PBS and lysed for 5 min on ice in lysis buffer (20 mM Tris-HCl (pH 7.4), 150 mM NaCl, 100 mM NaF, 10 mM Na_4_P_2_O_7_ and 10 mM EDTA supplemented with 1% (v/v) Triton X-100 and one complete protease inhibitor cocktail tablet (Roche)). After scraping, cells were centrifuged at 20,000g at 4°C for 10 min to remove cell debris and the lysis reaction was terminated by addition of 5x SDS loading buffer (250 mM Tris-HCl (pH 6.8), 10% (w/v) SDS, 40% (v/v) glycerol, 0.1% (w/v) bromophenol blue and 125 mM EDTA (pH 8.0)) supplemented with 10% β-mercaptoethanol. Samples were boiled for 5 min at 95°C. The proteins were separated on a NuPAGE 8.5% Bis-Tris Gel (Invitrogen) and transferred to a polyvinylidene difluoride (PVDF) membrane (Immobilon FL, Millipore) via a semi-wet process with the membrane first soaked in MeOH and then in transfer buffer (39 mM glycine, 48 mM Tris, 0.04% (w/v) SDS and 20% (v/v) MeOH). After the transfer, the membrane was incubated in blocking buffer (LI-COR) for 1 h and then for 4 h at R/T with the appropriate primary antibodies diluted in the blocking buffer at either 1:1000 dilution (pan Akt, pThr 308), 1:500 (γThr 514) or 1:1600 (α-Tubulin). After a first wash of the membrane (PBS with 1% (w/v) milk and 0.2% (v/v) Tween-20) to remove unspecific binding it was incubated for 1 h with the infrared dye-conjugated secondary antibodies (LI-COR) at a 1:5000 dilution each. The membrane was scanned on the Odyssey infrared imaging system (LI-COR) after a final wash.

### Quantitative IR Western blots

Western blotting using fluorescent IR secondary antibodies allows two antibodies from different origin to be detected simultaneously. We used a pan antibody (pan Akt, α-Tubulin) and the phospho-antibody indicated in each Figure. The fluorescent signal registered for each phospho-antibody was normalised to its corresponding pan antibody after background subtraction.

Phosphorylation of SGK1 of its T-loop at Thr 256 was quantified using the phospho-PKC(pan) antibody, which has been previously shown to recognise phosphorylation at the T-loop of several AGC kinases including SGK isoforms, S6K and PDK1 (Fig. S8) [41, 42]. The fluorescent signal was normalised to α-tubulin as indicated above.

### Protein purification

The DNA sequences coding for the fluorescently tagged species of PH^PDK1^ were cloned in pRSET-6xHisTag-TEV vectors and purified by affinity chromatography. The pRSET plasmids were transformed in BL21 cells. The TEV site was included in the sequence to cleave the His tag after the initial purification using a TEV protease. A final polishing process was done by size-exclusion chromatography.

### Protein lipid overlay assay

Commercial membrane lipid strips were purchased from Echelon (Salt Lake City, UT, USA). Otherwise, serial dilutions of PtdSer were spotted onto nitrocellulose membranes. In either case, the membranes were blocked for 1 h with a 3% BSA solution in TBS and they were incubated in 10 ml of a 30 μg/ml solution of recombinant eGFP-PH^PDK1^ species. Finally, the membranes were imaged using the fluorescent emission of eGFP to quantify the lipid-bound protein.

## Imaging

### Scanning Fluorescence Correlation Spectroscopy

Scanning FCS was applied to study the diffusional behaviour and intermolecular interactions of fluorescently-labelled proteins bound to GUVs (Fig. S2A). A water immersion objective HCX PL APO 63x/1.20 W CORR Lbd Bl (Leica) on a confocal Leica SP5 was employed and confocal detection of the emission using the APDs on the DD X1 port of the SP5. To excite eGFP tagged PH^PDK1^ species we used the 488 nm line from an Ar^+^ laser. The average power density at the sample plane was 50 kW/cm^2^. The filter for eGFP emission was a 500-530 nm bandpass. Photon arrival times were registered using an SPC-830 TCSPC card, which also registered the pixel, line and frame signals from the scanner. An HRT-82 Eight Channel Router for APDs (B&H) was required to spectrally tag and to invert the voltage of the photo-electron signals. An external clock signal at 20 MHz was used (Fig. S2B)

The imaging mode at the microscope was set to *xt* (line-scanning) and the scanning frequency was to 1400 Hz. The scanning length spanned over 16.64 μm, binned in 64 pixels of 260 nm each (x15 zoom). The membrane of every GUV was scanned for 5 min. The Becker & Hickl files were decoded using an in-house Matlab routine. The *xt* photon trace was aligned to correct for the vesicle drift (Fig. S2C) and the resulting photon time trace was fit to a 2D diffusion model with a minimal lag-time given by the inverse of the scanning frequency:

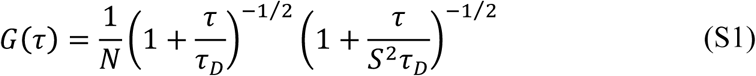

where N is the average number of molecules in the focal volume, *τ* is the lag-time, *τ_D_* is the diffusion time of the molecule and *S* ratio of the axial (*z*_o_) to the radial (beam waist, *ω*_o_) 1/e^2^ dimensions of the confocal volume. *ω*_o_ and *z*_o_ were obtained after calibrating the volume using a dye with known diffusion (Alexa 488, *D* ≈ 430 μm^2^/s at 25°C). Finally, the diffusion coefficient is *D* = *ω*_o_^2^/(4τ_D_).

### Confocal and time-resolved imaging

Confocal and lifetime images were acquired on a Leica TCS SP5 confocal scanning microscope through a HCX PL APO 63x/1.30 GLYC CORR CS 21°C objective (276 nm pixel size).

Lifetime imaging was performed using 100 fs pulses from an 80 MHz rep-rate Ti:Sapphire laser (Mai-Tai, Spectra-Physics) tuned at 890 nm, with an average power at the sample of 1 mW. The emission in the 500-550 nm range was registered on a hybrid detector on non-descanned configuration using a TCSPC system (Becker & Hickl SPC-830; IRF 170 ps FWHM).

Confocal steady-state intensity images were acquired at 488 nm and 543 nm He:Ne, respectively, right after IR-excited lifetime imaging and after correcting the focal plane for chromatic aberration. The emission was detected at 500-550 nm and 575-750 nm. The axial separation between the IR and the visible focal planes was calibrated upon IR and visible image registration at different planes, which yielded an average 1.2 μ separation between focal planes for the 63x/1.20W objective. The read-out for the fluorescence intensity was normalised to the excitation power and number of pixels in the region of interest of the image (Fluorescent Units). The average power density at the sample using CW visible lasers was 20 kW/cm^2^.

Live cell imaging was performed on a confocal configuration using a HCX PL APO 63x/1.20 W CORR Lbd Bl objective at 68 nm pixel-size and setting the scanning speed at 8000 Hz (resonant scanning). During image acquisition cells were kept in a 5% CO_2_ chamber at 37°C in observation medium (48 ml DMEM no phenol red (high glucose, no glutamine), 500 μl 100 mM Sodium Pyruvate, 500 μl of GlutaMax Supplement and 1 ml 10% BSA).

### Preparation of fixed samples for time-resolved FRET experiments

Cells were seeded at 300,000 per MatTek dish and transfected as explained above. After the 20 h of serum starvation period SKBR3 cells were stimulated with EGF to a final concentration of 100 ng/ml for 2 min. Non-starved NIH3T3 were stimulated with PDGF (30 ng/ml) for 2 min. PI3K inhibition was done prior to growth-factor stimulation treating the cells with 50 μM LY294002 for 20 min at 37°C and 5 or 10% CO_2_, depending on the cell line. During growth factor stimulation cells were maintained at the same conditions, respectively. Immediately after, cells were twice washed in 2 ml cold phosphate-buffered saline (PBS) and fixed in 1 ml PBS containing 4% paraformaldehyde for 12 min at R/T. SKBR3 were washed again with PBS and incubated at R/T for 10 min in 2 ml of a solution of sodium borohydride in PBS at 0.1% (w/v) to minimize the autofluorescence from the cells. Dishes were washed again with 2 ml of cold PBS two or three times until the bubbles from the sodium borohydride solution disappeared. After completely removing the remaining PBS, the dishes were mounted with circular coverslips (#1.5, Menzel-Glässer) using 20 μl of Mowiol mounting medium (10% (w/v) Mowiol 4-88 in 25% (v/v) water, 25% (v/v) glycerol and 200 mM Tris-HCl (pH 8.5)) containing 2.5% (w/v) 1,4-diazabicyclo(2.2.2)octane (DABCO), an anti-fade reagent.

### Image segmentation and calculation of the plasma membrane to cytoplasm partition coefficient

The confocal (visible excitation) and time-resolved (IR excitation) image of every cell was segmented into plasma membrane, cytoplasm and nucleus based on the confocal intensity images. The plasma membrane pixel-width was calibrated for every cell line and microscope resolution used on samples transfected with a membrane-bound probe (mRFP-Lyn or eGFP-PH^PDK1^). This figure was later used to outline the membrane of cells with poor membrane to cytoplasm intensity contrast. All measurements were normalised to the number of pixels in each region to account for the effect of the different area of every region. Morphological effects were this way reduced and the membrane to cytoplasm partition coefficient was empirically found to be constant within roughly one micron of the equatorial section (Fig. S4). Cells were always imaged in this region to avoid relative intensity variability at the polar regions.

### Quantification of accessible PtdIns(3,4,5)P_3_ and PtdSer

To monitor and quantify intracellular PtdIns(3,4,5)P_3_ we used the PH domain of GRP1, which binds PtdIns(3,4,5)P_3_ with high specificity [27]. Cells were transfected with μg of plasmid encoding a eGFP-GRP1^PH^ construct and allowed to express for at least 24 h after transfection. SKBR3 cells were either fixed or kept alive in an incubating chamber (37°C/ 5% CO_2_) after 20 h of serum-starvation. NIH3T3 cells were not serum-starved. Cells were imaged on the confocal microscope at 68 nm pixel size. Relative quantification of the PtdIns(3,4,5)P_3_ levels was performed by calculating the average fluorescence intensity of separated regions of the cell normalised to the cell-region pixel-area and the laser power (fluorescence units, FU) in a section near the equatorial plane. The pixels corresponding to the PM, the cytoplasm and the nucleus were analysed separately by image segmentation. The ratio of the normalised intensity at the PM to that at the cytoplasm (PM/Cyt ratio) was used to quantify differences in the translocation level of the GRP1^PH^ construct between cells of the same line. To facilitate the comparison, the PM/Cyt ratio of each cell was normalised to the average PM/Cyt obtained with a mCherry-6xGly-eGFP tandem in each cell line, used to quantify the level without translocation. The nucleus was excluded from the analysis by segmentation.

To quantify the accessible endogenous PtdSer content at the inner leaflet of the PM we used the discoidin-like C2 domain of bovine Lactadherin (LactC2) as a genetically encoded fluorescent biosensor that binds accessible PtdSer with high selectivity [30]. Imaging, transfection and segmentation were performed analogously to GRP1^PH^ as above. Reliable quantification was achieved by co-expressing mCherry-LactC2 and free-diffusing eGFP at a constant ratio by means of a self-cleaving P2A peptide (mCherry-LactC2-P2A-eGFP plasmid) [31]. As free-diffusing eGFP distributed homogeneously in the cytoplasm, its intensity was proportional to the total amount of mCherry-LactC2 in the cell. For every condition, the normalised average intensity of mCherry-LactC2 measured at the PM was plotted against the normalised average intensity of eGFP in the cytoplasm for a large population of cells and fit to a straight line. The slope of this linear regression allowed reliable quantification of relative PtdSer content at the PM in different conditions (Fig. S7).

### Quantification of the Dimerisation Efficiency (*E*_D_)

The dimerisation efficiency, *E*_D_, was calculated on a per cell basis for more than 30 cells per condition. All data was analysed following the methodology developed in [23]. The fluorescent decay from cells transfected with eGFP-PDK1 or eGFP-PH^PDK1^ was majorly monoexponential with a small contribution from a second component due to eGFP photophysics [43, 44] and possibly, residual autofluorescence. The lifetimes 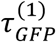 and 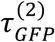 and the contribution of each component 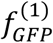 and 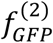 to the overall fluorescent emission are dependent on the molecular environment and fixation methods. Donor-only samples were first studied to characterise their fluorescent decay in our specific experimental conditions. Spatially invariant lifetime analysis of all cells transfected with donor-only species yielded 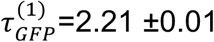 ns and 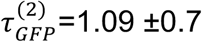 ns for SKBR3 cells and 2.16 ±0.01 and 0.77 ±0.05 ns for NIH3T3 cells. The contribution of the shortest lifetime 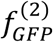 was 0.10 ±0.02 (N= 75 cells) for eGFP-PDK1 and eGFP-PH^PDK1^ irrespective of the cell line. No difference between mutants was found within the experimental uncertainty.

A spatially invariant analysis of the lifetime was performed on all images corresponding to a FRET pair (WT, mutant) and experimental condition (resting, Growth Factor-stimulated or PI3K-inhibited). It was assumed that only one molecular conformation for the interaction was possible (or, if they were more, they were indistinguishable within the experimental uncertainty). The fluorescent decays could be satisfactorily fit to a double exponential model with randomly distributed residuals (Fig. S3 A-B). The donor long lifetime of donor and acceptor co-transfected samples was similar to the long-lifetime of donor-only samples, 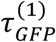, and was thus, identified as non-FRET donor [23]. The short lifetime was smaller than the short lifetime of donor-only samples 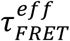, (0.87 ±0.03 ns for SKBR3 cells and 0.72 ±0.03 ns for NIH3T3 cells) and its corresponding weight, 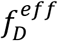, was higher. This component was, thus, identified as a mixture of donors involved in FRET and unperturbed donors relaxing with the fast-decaying component that had been previously identified in the absence of an acceptor [23]. In analogy to the simpler case of a monoexponentially decaying donor undergoing FRET, we termed this component an effective FRET component.

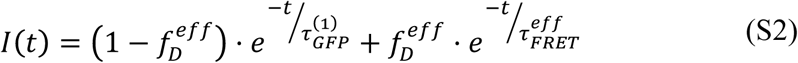

The 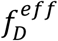 for the PDK1-PDK1 interaction was consistently different between mutants as well as before and after stimulation (Fig. S3C-D and S3A), confirming that the difference in FRET that we measured was due to a cellular response induced by the growth factor and not to random donor-acceptor encounters within the cell. Moreover, we experimentally ruled out non-specific FRET since the 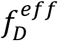 in the cytoplasm of cells that had been co-transfected with eGFP-PH^PDK1^ and free-diffusing RFP was at the lower detection limit except for cells with the highest concentration of acceptor. In all cases it was below that of cells that had been co-transfected with eGFP-PH^PDK1^ and mRFP-PH^PDK1^.

For a low affinity molecular interaction, as is PDK1-PDK1 homodimerisation [23], the slope of the linear regression of the fraction of bound donor as a function of the total acceptor concentration is roughly proportional to the affinity of the interaction. The total acceptor concentration, *A*_T_, is proportional to the fluorescence intensity of the acceptor. Therefore, under the low affinity assumption, the slope of the regression of 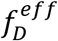 as a function of acceptor fluorescence provides an estimate of the probability of PDK1 homodimerisation. In the absence of acceptor, no FRET should occur and thus the intercept *A*_T_=0 is precisely the contribution of the fast-decaying component of eGFP, which had been also determined independently from donor-only samples 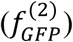.

However, the quantification of its uncertainty is challenging due to cell-to-cell variability and the fact that the linear regression is only an approximation to complex equilibrium behaviour. Consequently, rather than the linear regression, we used the mean of the distribution of 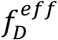 after correcting for the total intensity of the acceptor for every cell as an estimate of the probability of PDK1 homodimerisation, and its standard deviation as its uncertainty (Fig. S3E). This was, in turn, a measurement of the dimerisation efficiency (*E*_D_) of PDK1.

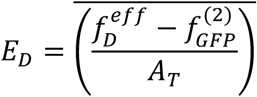

PDK1-Akt/PKB interaction was quantified in an analogous fashion after imaging the FRET between eGFP-PDK1 and mCherry-Akt/PKB.

All data concerning TCSPC fitting, image registering, and segmentation was analysed using in-house developed software written in Matlab. TCSPC fitting algorithms performed iterative-reconvolution with the experimentally characterised IRF and accounted for low photon-count time-bins at the tail of the decay. The FLIMfit software tool developed at Imperial College London and TRI2 (Paul R. Barber, University of Oxford) were occasionally used to render FLIM images.

### Segmentation of FRET-FLIM images

The *E*_D_ was first quantified for the whole cell. The image of every cell was afterwards segmented into plasma membrane, cytoplasm and nucleus based on the confocal intensity images (Fig. S3F) as explained above. The lifetime data for the cellular regions was fitted to a double exponential model fixing the long lifetime obtained from the spatially invariant analysis above. This way the *E*_D_ for the cellular regions was obtained.

### Molecular modelling

All experiments were performed with 1W1D [5] structure retrieved from the Protein Data Bank (PDB). This structure of the PH domain was used because of its excellent resolution (1.5Å), and the presence of inositol-(1,3,4,5)-tetrakis phosphate (Ins(1,3,4,5)P_4_) bound to the protein along with a glycerol molecule, suggesting binding sites. The N-terminal part of this structure corresponds to the linker between the PH domain and the kinase part (absent here). Accelrys Discovery Studio 3.1 was used to prepare the protein. Ligand ions, water molecules were removed manually and the resulting file subjected to the ‘Clean Protein’ module which allowed completion of all missing atoms or residues automatically. In this case, missing atoms were on the extremities and completed for the sake of coherence. The serine residue at position 410 was phosphorylated manually. After verifying the modifications and selecting alternate residues in order to minimise any steric hindrance, the protein was submitted to a cycle of minimisations, first with a Steepest Descent (SD) then a Conjugate Gradients algorithm with constraints on the backbone. Glycerol (GLY) and Ins(1,3,4,5)P_4_ were bound in the crystal at different sites which are biologically relevant [6]. During all the minimisation phases, both small molecules were considered as ligands and conserved then saved in different files before simulations. In order to check the viability of the phosphorylated protein, the obtained file was submitted to an equilibration simulation by molecular dynamics using GROMACS 5.0 package and the gromos 54a7 forcefield [45]. Protein was solvated with explicit water molecules and a 150 mM concentration of salt. The system was equilibrated in the constant-NVT (number of particles, volume and temperature) ensemble for 2 ns then in the constant-NPT (number of particles, pressure and temperature) ensemble for the same duration. For equilibrium purposes, positional restrains were applied on the protein backbone. Temperature and pressure were maintained at values of 300 K and 1 bar respectively. Since nothing is known about the phosphorylated PH domain, this calculation was performed to check its stability over 50 ns, asserting the possibility to use the 1W1D conformation, and the low RMSD obtained proved this assessment. During the rest of the simulations presented here, this conformation was kept rigid.

PtdSer was prepared after extraction from the PDB 3KAA structure, from which caproyl esters were removed [46]. PtdIns(3,4,5)P_3_ was prepared from the Ins(1,3,4,5)P_4_ molecule by adding a glycerol to the phosphate at position 1. Hydrogen atoms were added to all ligands, which were minimised briefly (SD algorithm) since all torsions were allowed during simulations. Vina 1.1.2 was used to dock small molecules, with full flexibility, while the receptor protein was kept rigid, as stated above. AutoDock Tools 1.5.6 were used to prepare the pdbqt files corresponding to the ligands and the receptor. Note that the fatty acid chains that should be present on both ligands were not considered here for the sake of simplification. Docking a ligand with two long carbon chains would not give accurate results in terms of geometry while lengthening considerably the calculation time. Once the ligand docked onto PDK1 surface, one has to view the fatty acid chains as perpendicular to the protein, embedded into a bilayer cellular membrane.

Three series of docking were obtained on the ‘GLY site’, in the ‘inositol site’ (site dockings) and on the whole surface of the receptor (blind docking) to assess any determinism. The respective boxes were used to define the docking site, taking into account the bound surface cavities with a 5Å margin in the first two cases, encompassing the whole receptor for the latter. This margin was reduced where the ‘GLY site’ and the ‘inositol site’ intersected to avoid encompassing the neighbouring cavity. For each docking site, five experiments were performed and analysed visually. The best poses (out of the maximum 20 obtained), those with the best energy were considered when they were identical over five simulations, leading to pose convergence. The following poses were generally convergent over the five experiments, but with slight energy differences. The ‘GLY site’ proved to be irrelevant in our experiments. Since a reasonable margin was used, all solutions were bound around the cavity previously occupied by the glycerol molecule, not inside it as determined in the original crystal. Even glycerol was not found preferentially in its native conformation, probably meaning its presence was a crystallisation artefact. The ‘inositol site’ allowed binding of the inositol “ligands” with conformations nearly identical to the one observed in the crystal structure (RMSD<1Å). PtdSer also bound to this site, not exactly in the cavity but slightly on one side, occupying the site previously held by one of the inositol phosphates, and forbidding any more docking into this region.

When blind docked, the “ligands” showed preferences, of the utmost importance. Glycerol did not bind into the inositol or in the ‘GLY site’, probably confirming its lack of relevance as a ligand in the native crystal. It did not show any selectivity but was bound in nearly all cavities of the protein surface. The Ins(1,3,4,5)P_4_ “ligand” bound the ‘inositol site’ albeit in a conformation slightly different from the crystal, while interacting with the known protein residues. On the contrary PtdIns(3,4,5)P_3_ and PtdSer specifically bound to the ‘inositol site’ with a high selectivity. When docked this way, binding of the inositols simultaneously with PtdSer was thus impossible, which implied a competition for the same binding site between these “ligands”. The energy of the docked ligands could be used to discriminate them, but considering the structural differences, and the small number of molecules this would seem abusive and was not the purpose of this study. None of the “ligands” specifically docked into the ‘GLY site’ in this experiment. One should note that the blind docking provided qualitative results about a ligand tendency to bind a specific area of the receptor.

In order to propose dimer formation, protein-protein docking was performed with the Patchdock server available on the Internet [47], using the automated procedure with no restrictions or preferences. Three PDK1 PH-domain structures were used in order to obtain complexes, where the proteins were rigid during the whole process. The first one was the one described above, that was used for docking small molecules without ligands in the active sites suggested in the crystal. The second one incorporated PtdIns(3,4,5)P_3_ in the ‘inositol site’, as obtained by site docking (note that the pose was nearly identical to the crystal). The last one showed the PtdSer and was obtained by docking as described above. These three structures were combined with Patchdock in order to obtain all six possibilities and simulate dimerisation of PDK1 PH domain with or without ligands. Among the obtained solutions, the 10 best ones were saved and examined visually, while Patchdock scoring was used to discriminate these. Three dimers were found with the highest scores among the 6 series, and compared by superimposition in order to identify these precisely with respect to the ‘inositol sites’ occupation. This means that one complex was found four times with different ligands. The presence of ligands in the ‘inositol site’ and a possible association with a membrane were checked to detect any structure hindrance forbidding the ligands to bind the lipid bilayer. This way, C1 and C2 complexes were recognised as able to bind a membrane through its ligand(s) in the ‘inositol site(s)’. On the contrary C3 displayed steric hindrance forbidding an access of PtdIns(3,4,5)P_3_ to a lipid bilayer. The angles measured between the PH domain in the dimers were obtained through the linkers direction after docking. This provided information about the proximity between the respective kinase domains in the obtained dimer.

## Supplementary Figures

- Fig. S1: Fluorescently-tagged PDK1, PH^PDK1^ and Akt/PKB constructs.
- Fig. S2: Experimental implementation of scanning FCS.
- Fig. S3: Quantification of the binding efficiency of two proteins using Time-Resolved FRET (FRET-FLIM).
- Fig. S4: Calculation of the plasma membrane to cytoplasm partition coefficient.
- Fig. S5: Quantification of PtdIns(3,4,5)P_3_ levels in intact NIH3T3 and SKBR3 cells using the high-selectivity PH domain probe eGFP-GRP1^PH^.
- Fig. S6: Time course of PtdIns(3,4,5)P_3_ levels at the plasma membrane in live NIH3T3 and SKBR3 cells using eGFP-GRP1^PH^.
- Fig. S7: Quantification of PtdSer levels at the plasma membrane of NIH3T3 and SKBR3 cells.
- Fig. S8: Use of phospho-PKC(pan) (γThr 514) to detect phosphorylation of endogenous SGK1 at its T-loop.

**Fig. S1:**
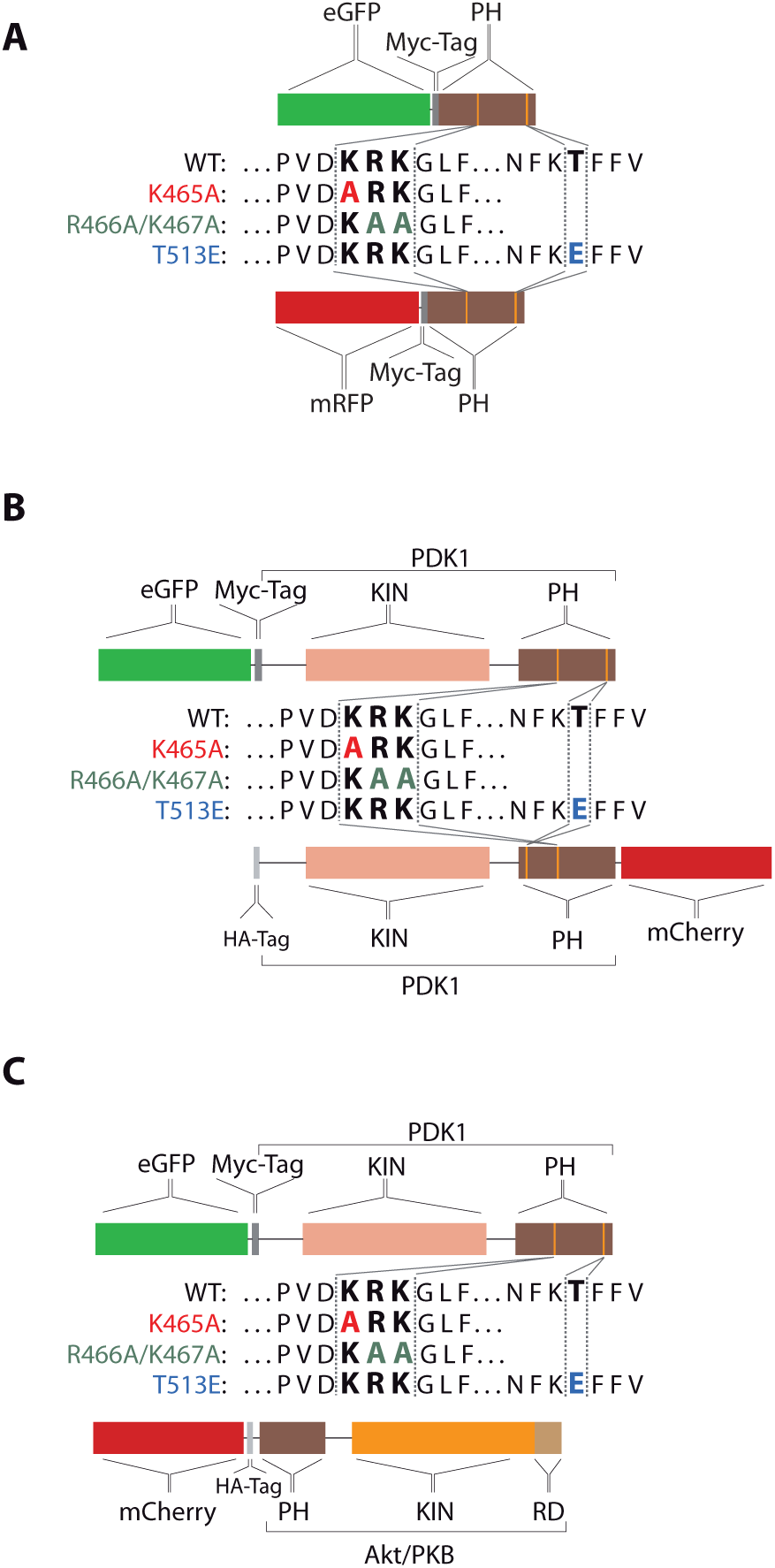
Fluorescently-tagged PDK1, PH^PDK1^ and Akt/PKB constructs.

**Fig. S2:**
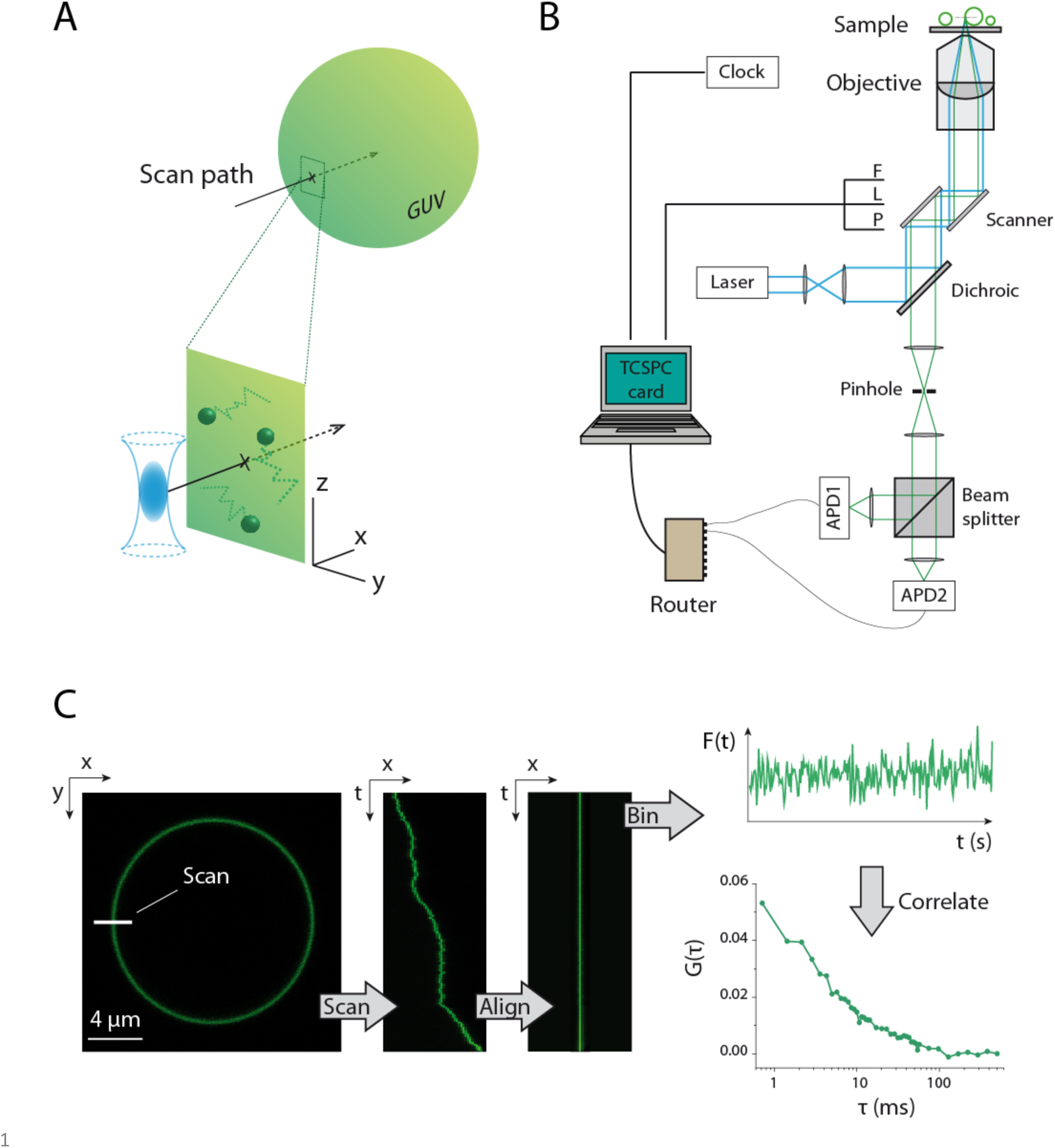
Experimental implementation of scanning FCS. **(A).** Scanning FCS in GUVs was performed along a path perpendicular to the membrane plane (yz) and to the optical axis (z). **(B)** Instrumental configuration for point and scanning FCS, explained in the text. F, L and P refer to the frame, line and pixel signals sent by the microscope to the TCSPC card in order to track the scanner position in the sample. **(C).** The GUV was placed near its equatorial plane and a single line was scanned for 5 min at 1400 Hz. Due to the GUV drift the position of the membrane in the scanned line was shifted during the measurement. Rectification of the membrane position in time was done by alignment of the fluorescence maxima in each line. Spatial binning of the pixels corresponding to the GUV membrane allowed obtaining the fluorescence fluctuation time trace, which was used to calculate the ACF. The maximal temporal resolution of this ACF was limited by the scanning frequency.

**Fig. S3.**
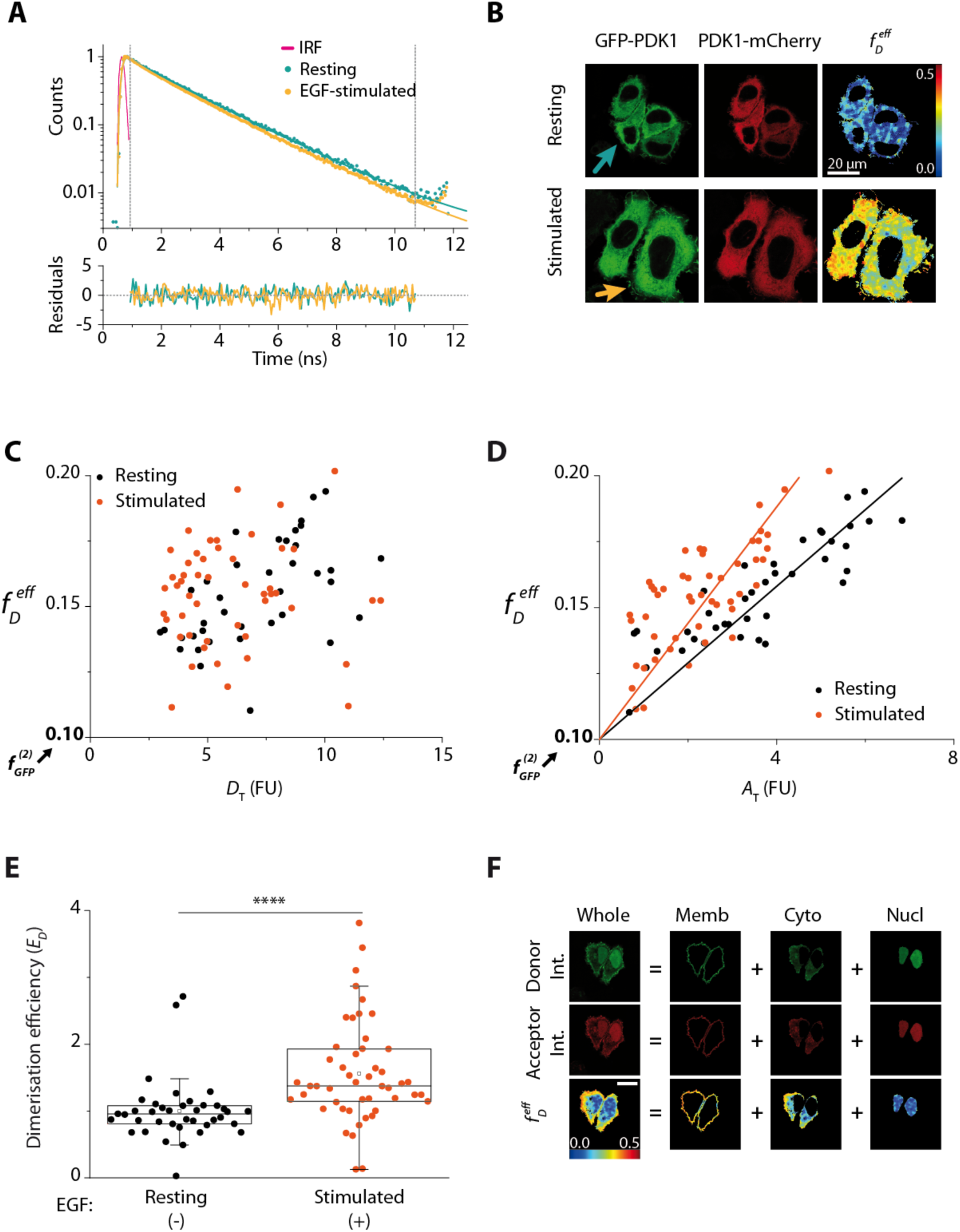
Quantification of the binding efficiency of two proteins using Time-Resolved FRET (FRET-FLIM). **(A-B)** Cells were transfected with proteins tagged to a fluorescent donor or acceptor. The effective fraction of donor undergoing FRET, 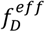, was calculated fitting the fluorescence decay to a biexponential model keeping the two lifetimes fixed to those obtained by spatially invariant analysis. **(A)** Representative decays of cells in resting (blue arrow) and growth factor stimulated conditions (yellow arrow): cell in resting conditions: 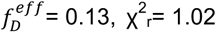; stimulated cell: 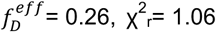. **(C-D)** The fraction of interacting donor depends on the donor and acceptor concentration expressed in the cell (each dot is one cell), measured as intensity normalised to the cell pixel-area and the laser power (FU, Fluorescence Units). The *f* D was found to be consistently higher after stimulation, ruling out that FRET is due to random-acceptor encounters. **(D)** Linear regression of 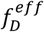 as a function of the acceptor concentration for different physiological conditions. **(E)** Binding efficiency for the donor-tagged protein quantified from (D) for resting versus EGF-stimulated cells under the assumption of a low affinity interaction. **(F)** The image of every cell was segmented into plasma membrane, cytoplasm and nucleus based on the confocal intensity images and the lifetime was calculated for every region.

**Fig. S4.**
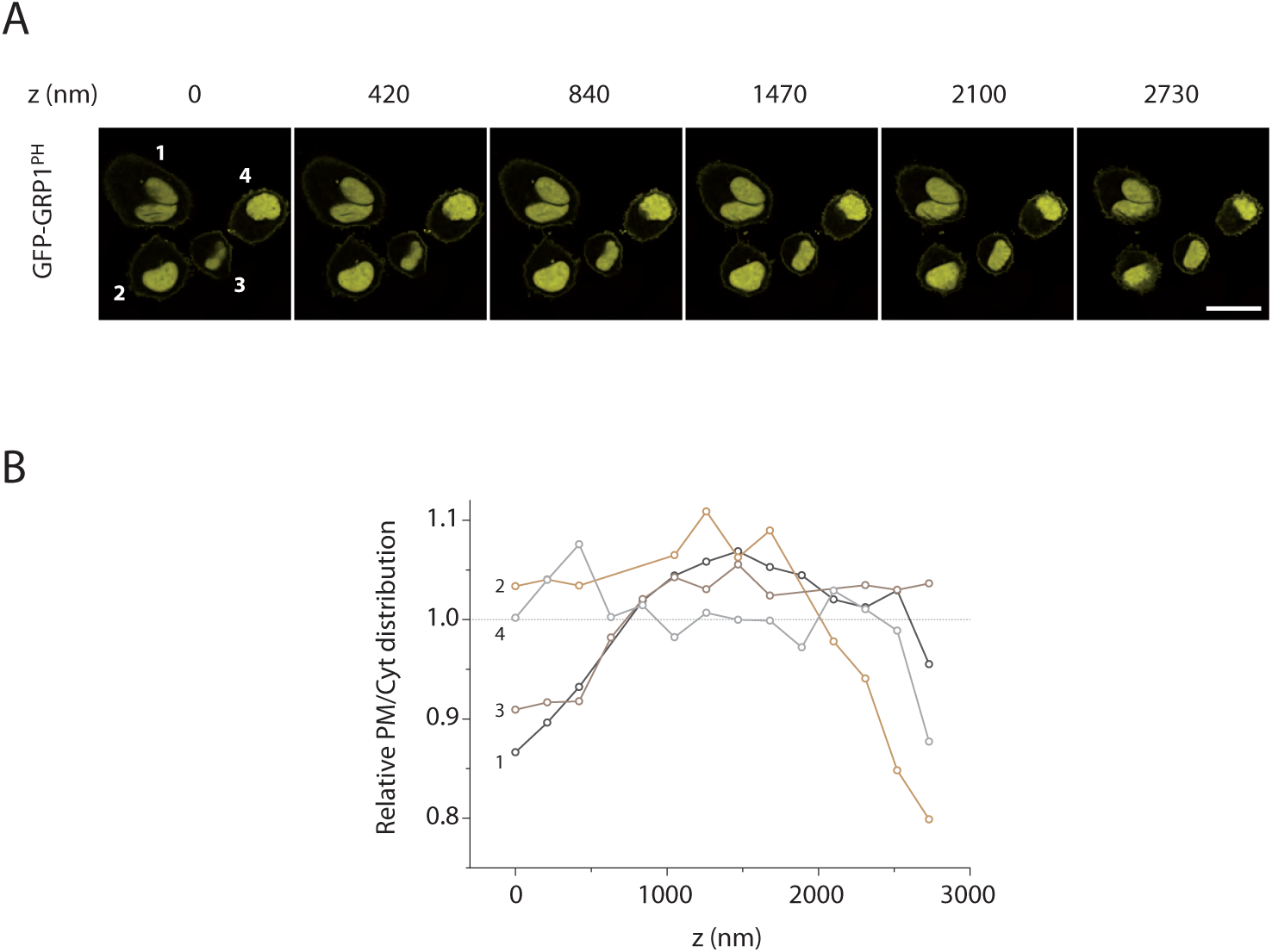
Calculation of the plasma membrane to cytoplasm partition coefficient. It is independent of the axial plane around the equatorial plane of the cell. **(A)** Representative z-stack of SKBR3 cells expressing GFP-PH^PDK1^. **(B)** The PM to cytoplasm ratio of the intensity of GFP-PH^PDK1^ was calculated for each axial plane and normalised to the average of each cell. The PM/Cytoplasm intensity ratio was found constant within roughly one micron of the equatorial section. Cells were thus imaged in this region to avoid variability. Scale bar: 20 μm.

**Fig. S5:**
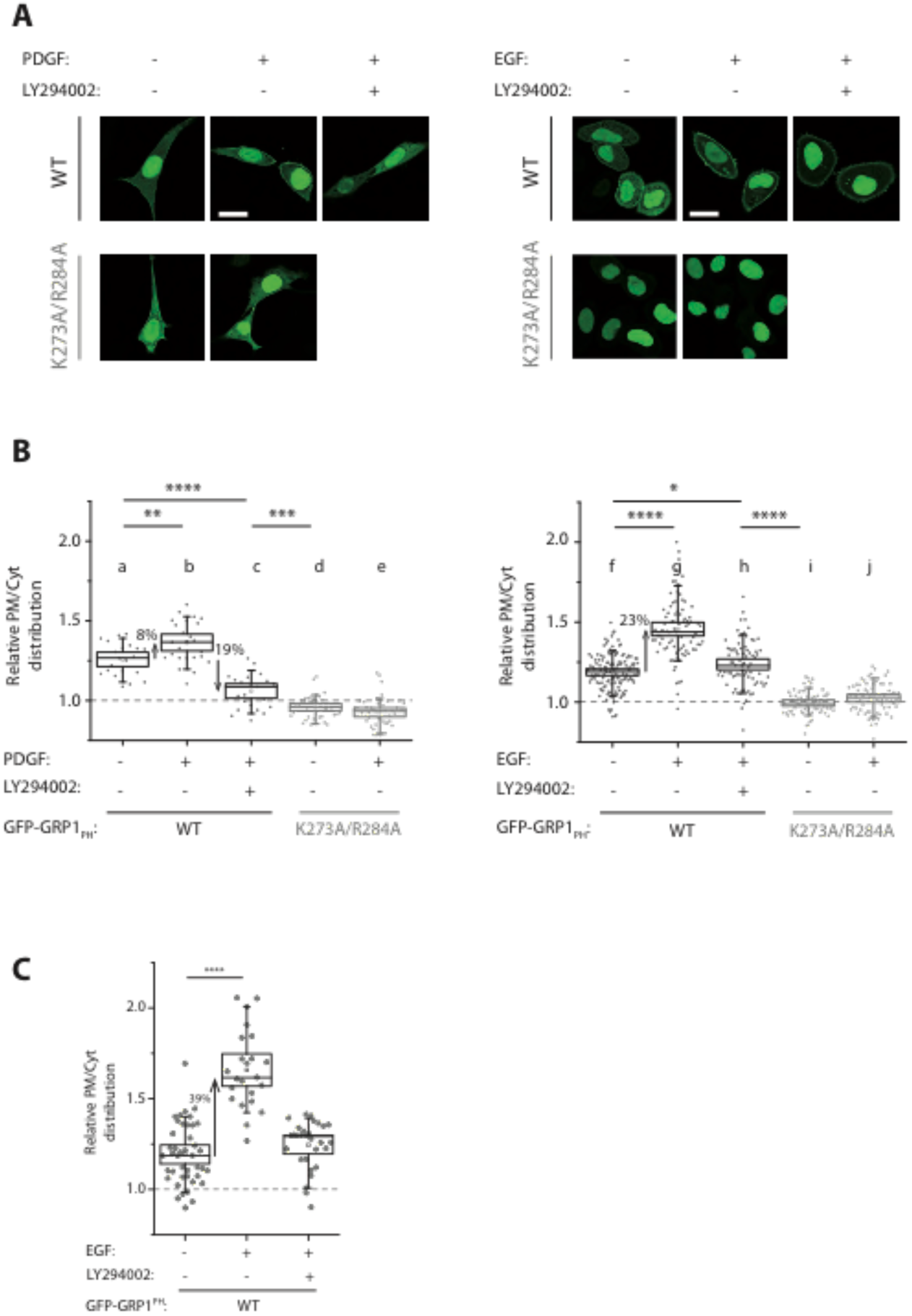
Quantification of PtdIns(3,4,5)P_3_ levels in intact NIH3T3 and SKBR3 cells using the high-selectivity PH domain probe GFP-GRP1^PH^. Left: NIH3T3 cells; Right: SKBR3 cells. All experiments were performed in resting conditions (-), growth-factor stimulated (+) and pretreatment with PI3K inhibitor LY294002 prior to stimulation. **(A)** Representative images of NIH3T3 and SKBR3 cells cotransfected with GFP-GRP1^PH^. No association of the non-PtdIns(3,4,5)P_3_-binding mutant (K273A/R284A) of GFP-GRP1^PH^ to the PM could be observed. **(B)** The amount of GRP1^PH^ accessible to PtdIns(3,4,5)P_3_ at the PM upon stimulation of the PI3K pathway is higher for SKBR3 cells than for NIH3T3 cells. When PI3K is inhibited prior to stimulation the amount of PtdIns(3,4,5)P_3_ at the PM falls below the basal levels in NIH3T3 cells, but stays at basal level in SKBR3. Scale bar: 20 μm. NIH3T3: N>20. SKBR3: N>30. Box: 2xSEM (95.4% confidence); Whiskers: 80% population. Mann-Whitney test *p<0.05. Three independent experiments. **(C)** Non-starved SKBR3 cells transfected with GFP-GRP1^PH^ were fixed and imaged in resting conditions, EGF-stimulated and treated with 50 μM LY294002 prior to stimulation to prevent PtdIns(3,4,5)P_3_ generation by PI3K. The distribution of GFP-GRP1^PH^ at the PM relative to the cytoplasm is analogous to starved cells for the three conditions. This result confirms that the lack of decrease of the PtdIns(3,4,5)P_3_ levels below basal in SKBR3 cells when PI3K is inhibited, is not due to an already reduced basal level of PtdIns(3,4,5)P_3_ due to starvation. Absolute values differ in each case. N>20.

**Fig. S6:**
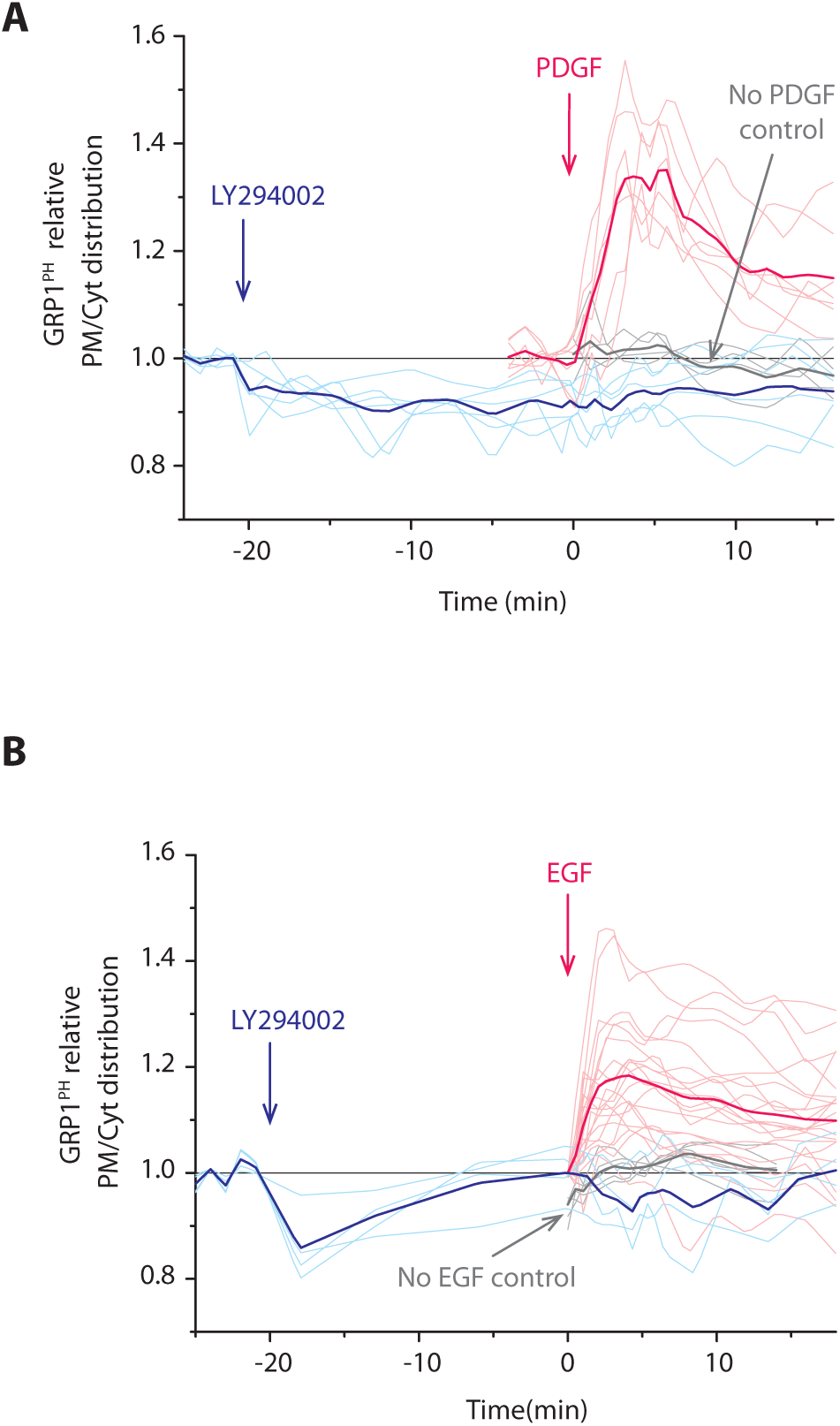
Time course of PtdIns(3,4,5)P_3_ levels at the plasma membrane in live NIH3T3 and SKBR3 cells using GFP-GRP1^PH^. PtdIns(3,4,5)P_3_ at the PM increases upon stimulation and is reduced below basal level after PI3K inhibition in live NIH3T3 (A), but not in SKBR3 (B) cells confirming observations in fixed cells. **(A)** NIH3T3 and **(B)** SKBR3 cells were transfected with GFP-GRP1^PH^ and imaged live at physiological conditions for more than 30 min after growth factor stimulation and/or inhibition of the PI3K pathway. The graph shows the distribution of the PH domain of GRP1 at the PM relative to the cytoplasm. The pink curves are cells stimulated with PDGF (NIH3T3) or EGF (SKBR3) at time 0; the blue ones are cells that had been treated with LY294002 for 20 min prior to stimulation at time 0; the grey curves are cells that had not been stimulated nor treated with the inhibitor. Highlighted curves are the average of the dimmer curves (NIH3T3: N_PDGF_= 7; N_LY_= 6; N_CTL_= 4; SKBR3: N_EGF_= 21; N_LY_= 4; N_CTL_= 5). The temporal distribution of single cells confirms the ensemble observations in fixed cells, shown in the main text.

**Fig. S7.**
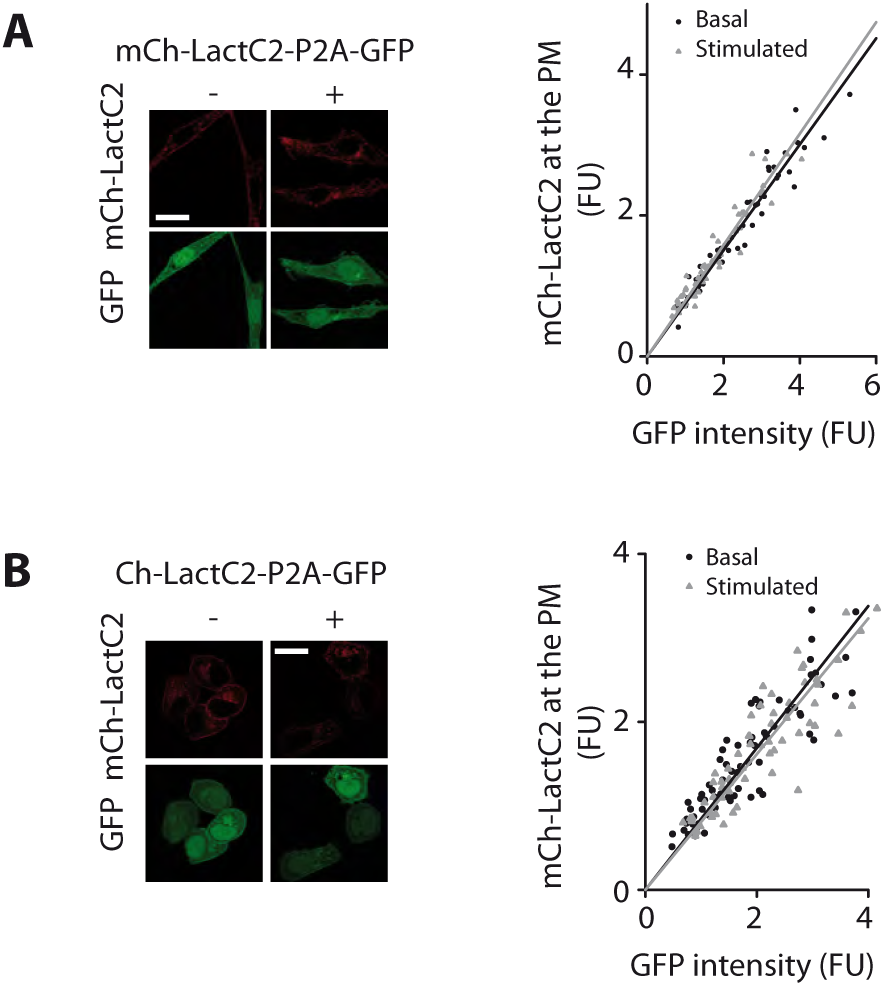
Quantification of PtdSer levels at the PM of NIH3T3 and SKBR3 cells. **(A)** NIH3T3 and **(B)** SKBR3 cells were transfected with the multicistronic plasmid mCherry-LactC2-P2A-GFP, which translates mCherry-LactC2 and GFP separately. The images in the top row show the mCherry-LactC2 channel and the ones in the bottom row that of GFP. GFP serves as a transfection reporter and also allows LactC2 expression to be quantified due to the constant ratio of P2A translation. Cells on the right column were stimulated with PDGF (NIH3T3) or EGF (SKBR3) for 2 min prior to fixation. The average intensity of mCherry-LactC2 at the PM was plotted as a function of that of GFP over the whole cell for every cell and fitted to a linear model. Differences in PtdSer localisation at the PM were not observed before or after growth factor stimulation. Scale bar: 20 μm. N>30. Three independent experiments.

**Fig. S8:**
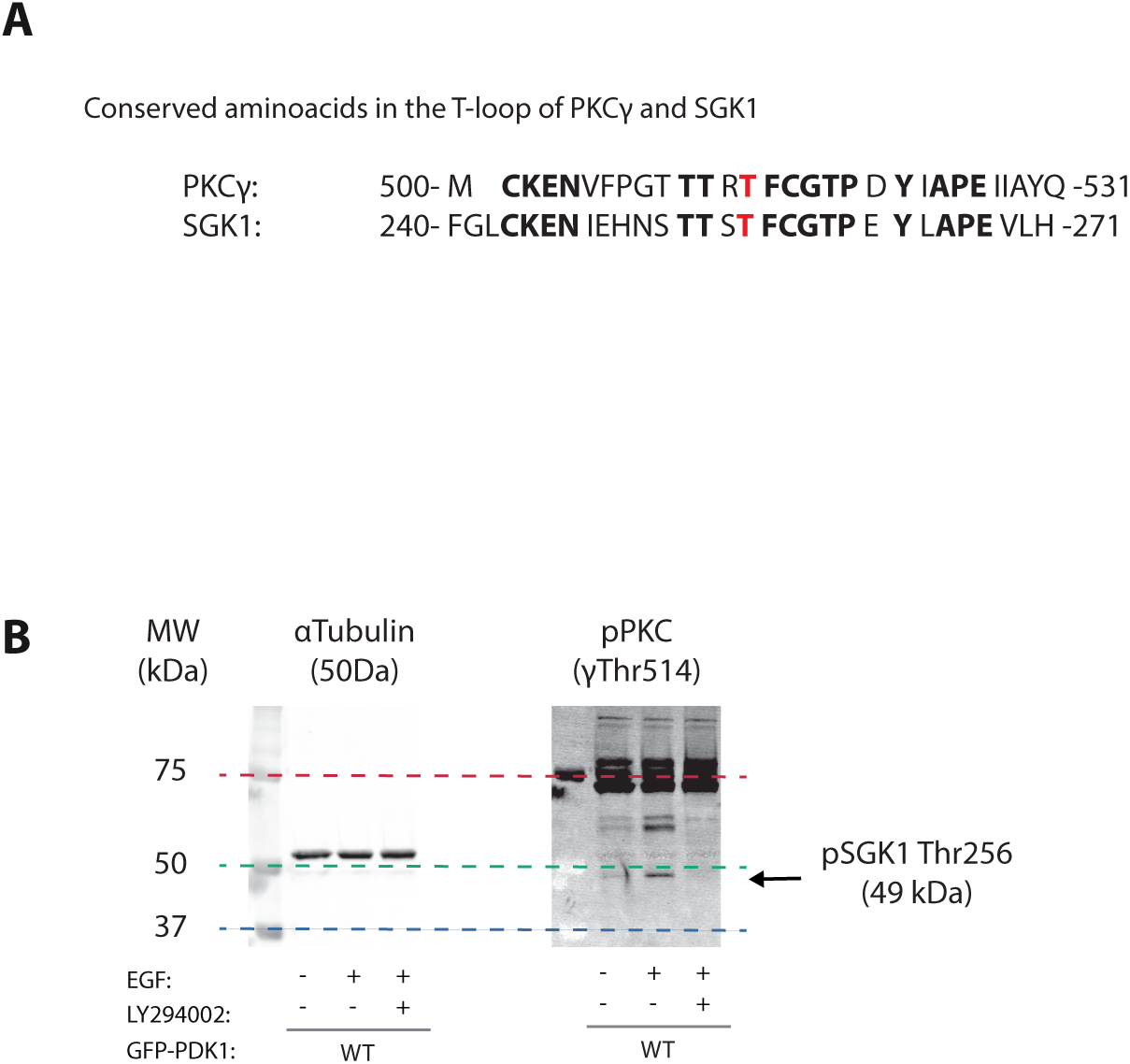
Phospho-PKC(pan) (γThr 514) detects phosphorylation of endogenous SGK1 at its T-loop. **(A)** The activation segment (T-loop) of PKC and SGK1, located in their catalytic domain, is a highly conserved regulatory motif. The phosphorylation residues for PKCγ (Thr 514) and SGK1 (Thr 256) are in red. **(B)** The phospho-PKC antibody efficiently recognises the T-loop of several AGC kinases, including SGK1 and PDK1. Endogenous phosphorylated SGK1 in Fig. 4D was quantified at the 49 kDa band (see Methods).

## Acknowledgements

We are grateful to Peter J Parker, Len Stephens, Patrick Williamson and Vytas Bankaitis for scientific discussions and constructively reading the paper. Sonia López Fernandez for her help in the cloning.

## Authors’ Contribution

G.dlH-M. performed the experiments and contributed to their design, V.C. contributed to the experimental design and co-wrote the manuscript; S.L.F. prepared the expression constructs and westerns; R.B. and J.D. performed the molecular modelling; B.L. and J.R.-I. directed the research, analysed the data and wrote the paper.

